# Therapeutic targeting of immunometabolism in Alzheimer’s disease reveals a critical reliance on Hexokinase 2 dosage on microglial activation and disease progression

**DOI:** 10.1101/2023.11.11.566270

**Authors:** Juan F. Codocedo, Claudia Mera-Reina, Peter Bor-Chian Lin, Shweta S. Puntambekar, Brad T. Casali, Nur Jury, Pablo Martinez, Cristian A. Lasagna-Reeves, Gary E. Landreth

**Affiliations:** Stark Neurosciences Research Institute, Indiana University, School of Medicine, Indianapolis, IN 46202, USA

**Keywords:** .: Microglia, Alzheimer’s disease, Hexokinase 2, amyloid, inflammation, Lonidamine, mitochondria, NFKβ

## Abstract

Microgliosis and neuroinflammation are prominent features of Alzheimer’s disease (AD). Disease-responsive microglia meet their increased energy demand by reprogramming metabolism, specifically, switching to favor glycolysis over oxidative phosphorylation. Thus, targeting of microglial immunometabolism might be of therapeutic benefit for treating AD, providing novel and often well understood immune pathways and their newly recognized actions in AD. We report that in the brains of 5xFAD mice and postmortem brains of AD patients, we found a significant increase in the levels of Hexokinase 2 (HK2), an enzyme that supports inflammatory responses by rapidly increasing glycolysis. Moreover, binding of HK2 to mitochondria has been reported to regulate inflammation by preventing mitochondrial dysfunction and NLRP3 inflammasome activation, suggesting that its inflammatory role extends beyond its glycolytic activity. Here we report, that HK2 antagonism selectively affects microglial phenotypes and disease progression in a gene-dose dependent manner. Paradoxically, complete loss of HK2 fails to improve AD progression by exacerbating inflammasome activity while its haploinsufficiency results in reduced pathology and improved cognition in the 5XFAD mice. We propose that the partial antagonism of HK2, is effective in slowed disease progression and inflammation through a non-metabolic mechanism associated with the modulation of NFKβ signaling, through its cytosolic target IKBα. The complete loss of HK2 affects additional inflammatory mechanisms associated to mitochondrial dysfunction.

**Highlights:**

- Hexokinase 2, the first and rate-limiting enzyme of glycolysis, is specifically upregulated in plaque-associated microglia of AD mice models and in the postmortem cortex of human AD patients.
- Microglia haploinsufficient in HK2 exhibit reduced amyloid burden and inflammation as well as improved cognition in a mouse model of AD. Paradoxically, the complete loss of HK2 results in opposite effects, by exacerbating inflammation.
- Lonidamine, an anticancer drug that inhibits HK2, mimics the salutary effects of HK2 haploinsufficiency in the 5xFAD mice, but only in males during the early stages of disease.
- HK2 deletion induced mitochondrial dysfunction associated to increased expression of inflammasome elements and IL-1β.
- HK2 partial antagonism exerts beneficial effects independent of its energetic or mitochondrial role, likely through cytosolic stabilization of IκBα and inhibition of the NF-κB pathway, leading to reduced proinflammatory gene expression.

## Introduction

Alzheimer’s disease (AD) is a highly prevalent neurodegenerative disorder and the most common cause of dementia in the elderly. AD pathology is characterized by amyloid β (Aβ) deposits and neurofibrillary tangles, both linked to neuronal death and cognitive decline. In the AD brain, a subset of microglia respond to amyloid deposition by migration to the nascent plaque where they proliferate^1^ and subsequently envelop the plaque with their processes, constituting a biological barrier segregating the plaque from the brain parenchyma^2^. The plaque-associated microglia undergo a dramatic morphological transformation that ultimately accompanies the acquisition of a proinflammatory phenotype, with the release of neurotoxic factors such as cytokines and an array of immune mediators. More recently, microglial activity has been associated with the seeding and spreading of new Aβ plaques^3^. and tau aggregates^4^. The microglial response to AD pathology is highly complex, the simultaneous exhibition of phenotypes associated with both disease exacerbation and mitigation that changes with disease progression has confounded the understanding of its mechanistic basis and the effects of potential therapies^5^.

The acquisition of the AD-associated phenotypes requires large amounts of energy to support the synthesis of cytokines, induction of chemotaxis, proliferation, and phagocytosis that accompany a significant increase in gene expression^6–8^. These energy-demanding responses lead to the reprogramming of microglial metabolism. For instance, homeostatic microglia employ mitochondrial oxidative phosphorylation to produce cellular ATP. In contrast, microglia respond to immune stimuli by undergoing a shift to aerobic glycolysis, which allows the rapid generation of ATP and other metabolic intermediates to power the various energy intensive activities that subserve the change in microglia phenotypes^9^. However, because glycolysis is metabolically inefficient, persistent reliance on glycolysis is postulated to lead to impaired immune function normally associated with chronic and exaggerated inflammatory responses^10^. During the last decade, several studies have described a plethora of mechanisms by which glycolysis orchestrates inflammation, including the buildup of metabolites that can regulate inflammatory gene expression. It has only recently been appreciated that glycolytic enzymes can regulate inflammation by the gain of non-metabolic functions that have gone undetected previously. These non-traditional mechanisms can be initiated by changes in the subcellular localization of glycolytic enzymes allowing them to bind and modify the activity of different inflammatory regulators^11^. These non-metabolic activities, add a new layer of regulation (complexity) to the relation between metabolism and inflammation.

Hexokinases perform the first and rate-limiting step in glycolysis through the irreversible phosphorylation of glucose to glucose-6-phosphate (G-6-P). The phosphorylated form of glucose is trapped inside the cell and is then further metabolized by glycolysis to generate ATP^12^. Like other glycolytic enzymes, HK2 has non-metabolic roles that can impact inflammation independently of its canonical glycolytic role. HK2 binds to the outer mitochondrial membrane (OMM) through direct interaction with the voltage-dependent anion channel (VDAC) and its displacement induces the formation of a mitochondrial pore that triggers the activation of NLRP3 inflammasome^13^.

Recent publications demonstrate the role of Hexokinase 2 (HK2) as a critical determinant of microglial metabolism and immune response in various pathological processes^14–16^. However, these independent studies reported conflicting data with respect to the effect of genetic inactivation of both alleles of the HK2 gene on microglial phenotypes and disease progression. Specifically, Leng, *et al.* described a reduction in inflammation and increased migration that attenuated disease progression in the 5xFAD mice^14^. In contrast, Hu *et al*, found a polar opposite response, whereby the complete deletion of HK2 induced an enhanced inflammatory response, reduced migration and increased brain damage in a murine model of cortical stroke^15^. The discrepancy is also mechanistic, as both groups present diametrically conflicting results in their *in vitro* assessment of ATP levels and use of alternative fuels by HK2 deficient microglia^14,15^. Thus, there is no consensus on the role of microglial HK2 in brain disease and its role in maladaptive inflammation, remains poorly understood.

In this study we investigated HK2 gene dosage in the regulation of microglial phenotypes and AD progression in the 5xFAD mice. Its inducible deletion modulates specific features of the microglial response in a gene dose dependent manner. In agreement with the report of Hu *et al*, the complete loss of HK2 results in a robust inflammatory phenotype that in the 5xFAD mice fails to alter Aβ plaque burden and attenuation of cognitive decline. Remarkably, HK2 haploinsufficiency results in very different effects, with attenuation of inflammation, reduction of Aβ plaque load and reversing disease-related cognitive decline. Importantly, we found that pharmacological inhibition of HK2 mimics the salutary effects of HK2 haploinsufficiency and partial loss of function in male 5xFAD mice, during the early stages of the disease, suggesting a common mechanism and that its targeting has therapeutic potential.

In our hands, the beneficial effects of the partial antagonism of HK2, was not associated with increased levels of ATP, lipoprotein lipase (LPL) up-regulation or mitochondrial disfunction as previously proposed^14^. Instead, we observed that the decreased catalytic activity of HK2 was associated with the increase in the levels of its cytosolic target IKBα, a regulatory protein that inhibits the nuclear translocation of NFKβ and the induction of the expression of inflammasome genes. In contrast, the complete deletion of HK2, overrode this mechanism by inducing mitochondrial dysfunction and inflammasome activation as previously described by Hu *et al* and others^13,15,17^.

## Results

### HK2 expression is induced in Aβ plaque-associated microglia

We first evaluated the expression of the predominant HK isoforms in the cortex of 5xFAD mice during disease progression. We found that the levels of HK2 were significantly increased in the 5xFAD mice compared to non-transgenic mice as early as 4 months and at 4-fold higher levels by 8 months of age (Fig. 1A-C). The changes in HK2 were significantly greater in female 5xFAD mice compared to males (Fig. 1A-C) which is consistent with female 5xFAD mice exhibiting higher levels of Aβ and augmented inflammation^18^. In contrast, the levels of the ubiquitously expressed HK1 were unchanged with disease progression (Fig 1B and C).

**Fig 1.**
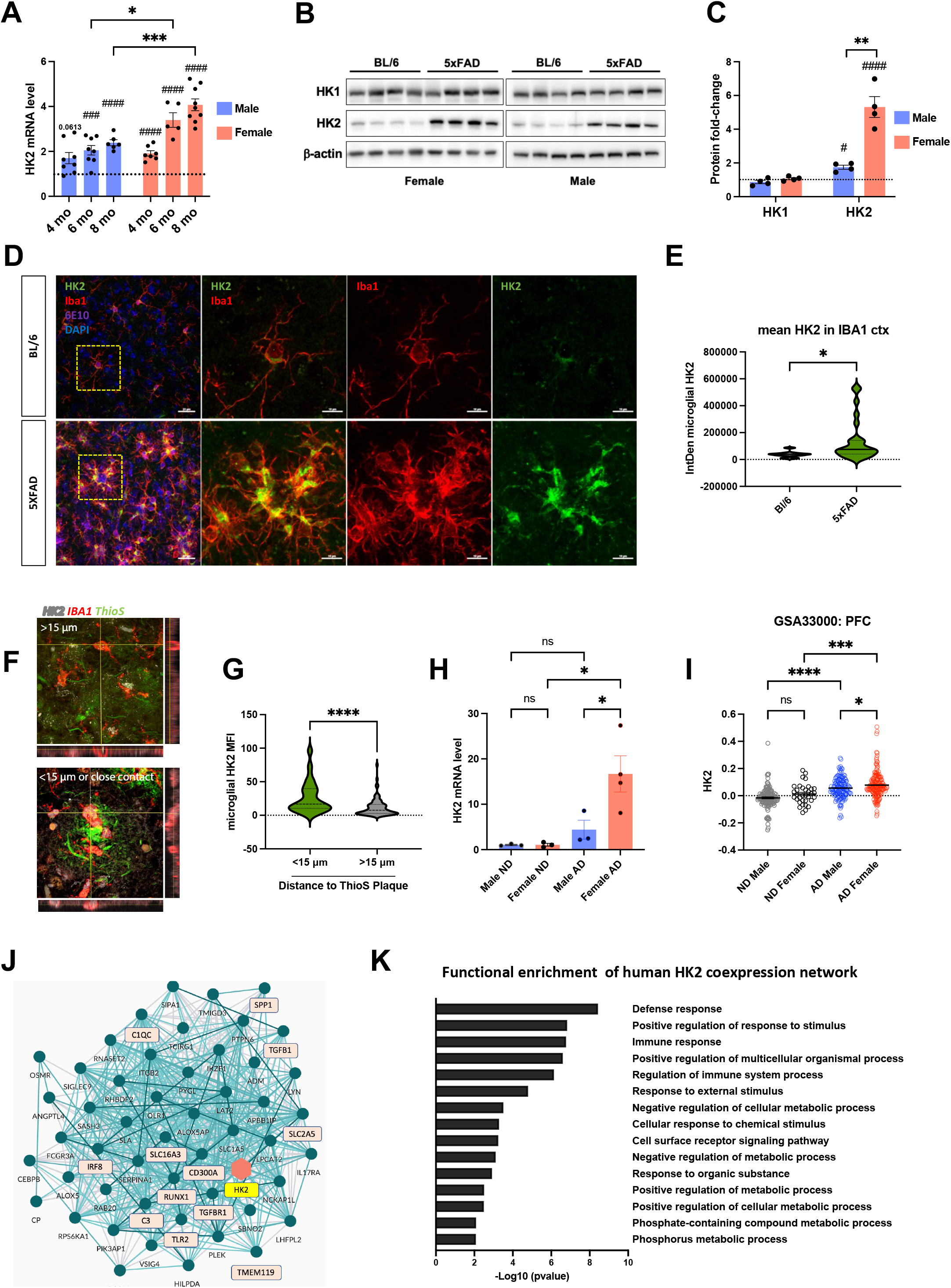
HK2 as a molecular hub between metabolism and immune response in AD. **(A)** qPCR analysis of HK2 expression during disease progression in the cortex of male and female 5xFAD mice. Dashed line represents gene expression in Bl/6 control mice. Post hoc analysis revealed significant genotype-related increases (###p< 0.001, ####p< 0.0001) and a significant increase in HK2 expression in the cortex of female, compared with male 5xFAD mice at 6 and 8 months (*p< 0.05, ***p< 0.001, n=5-9 per group, One-way ANOVA followed by Tukey’s post-hoc test). **(B-C)** Western blot and quantification of HK1 and HK2 in the cortex of 8-mo old 5xFAD mice. Dashed line represents gene expression in Bl/6 control mice. Densitometric analysis of hexokinases isoforms were normalized to GAPDH levels (n=4, genotype difference: #p< 0.05, ####p<0.0001 and sex difference: **p<0.01. Unpaired t test). **(D)** Confocal images of microglial HK2 expression in the cortex of 8-month-old BL/6 (upper panels) and 5xFAD mice (lower panels). Microglia (Iba1+, red), HK2 (green) and deposited amyloid (6E10, purple). Scale bar: 25 μm. **(E)** Quantification of the integrated density of HK2 in Iba1+ cells. After binarization of Iba1 immunoreactive signal, the mean fluorescence intensity of HK2 was calculated and multiplied by the area of Iba1. *p<0.05 (n=8-9 per group, Unpaired t test) **(F)** Immunofluorescent staining of HK2 (white), Iba1 (red) and ThioS (green) of brain samples of AD patients. **(G)** Quantifications of mean fluorescent intensity of microglial HK2 in close contact with Aβ (in radius <15μm form Aβ) and away from Aβ (>15μm). *p<0.05 (n=8-9 per group, Unpaired t test). **(H)** qPCR analysis of HK2 expression in female vs male post-mortem samples of non-demented (ND) and AD patients normalized to the expression of Iba1^+^ (n=3-4). **(I)** HK2 expression obtained from a human transcriptomic dataset of prefrontal cortex tissue (PFC) of 157 nondemented controls and 310 AD patients (GSE33000). Data segregated by sex reveals a significant difference in the up-regulated levels of HK2 between woman and men. **(J)** AMP-AD consortium co-expression network analysis of RNA-seq data from AD cases and controls. HK2 node, is highlighted in yellow and co-expressed genes associated to microglial activation or identity are highlighted in pink. The network analysis performed by the AMP-AD consortium, uses an ensemble methodology to identify genes that show similar co-expression across individuals. **(K)** Gene ontology of HK2 co-expression network. The list of specific microglial genes co-expressed with HK2 in hAD cases were used to evaluate the top 15 biological process associated with its expression.

To confirm that the increase in the levels of HK2 in the cortex are driven exclusively by microglia, we depleted microglia by treatment with the colony-stimulating factor 1 receptor (CSF1R) antagonist PLX5622^19^, which resulted in a complete loss of HK2 induction with no effect on HK1 (Fig. S1A and B). Furthermore, following inhibitor withdrawal and microglial repopulation, we observed a complete restoration of HK2 levels (Fig. S1A and B). By flow cytometry analysis, we observed a significant increase in the cellular expression of HK2 in microglia (CD11b^+^, CD45^+^) in 8-month-old 5xFAD mice compared to their non-transgenic controls (Fig. S1C and D).

To determine the identity of the microglial sub-population responsible for the HK2 increase, we performed immunohistochemical (IHC) studies of subiculum and cortex of 8-month-old 5xFAD mice and non-transgenic controls. In the BL/6 mice, staining with IBA1 revealed evenly tiled, ramified microglia, and these homeostatic cells, displayed faint somatic staining for HK2 (Fig. 1D, upper panels). In contrast, 5xFAD mice at 8 months of age exhibit clustered, plaque associated microglia that have an amoeboid morphology normally associated with the DAMs or MGnD phenotypes^20,21^. This subset of microglial cells exhibits increased immunofluorescence for HK2 throughout the microglial soma and processes that envelop the plaques (Fig 1D, lower panels and 1E). In fact, the regional increase in HK2 levels is directly associated to the amyloid load, being the subiculum the area with higher levels of HK2 (Fig. S1E and F). These observations suggest that induction of HK2 was triggered by direct contact of the microglia with deposited amyloid and could explain the time dependent induction of HK2 in the 5xFAD brain.

To determine whether these changes are conserved in human disease, we evaluated HK2 levels in the cortex of AD patients. As expected, we observed a significant increase in HK2 levels in the microglia in close association with Aβ-plaques (Fig.1F and G). Similar to our animal studies, qPCR analysis shows that the increase in HK2 levels was significantly greater in women compared to men (Fig. 1H).

To confirm our observation through independent databases, we evaluated the expression of HK2 in databases of gene expression in human AD from gene expression omnibus (GEO) and we found that the levels of HK2 are increased in the prefrontal cortex of a cohort of 310 of human patients. When we segregated the data by sex, the increase in women is significantly higher than men (Fig. 1I). Additionally, we observed that the levels of the neuronal isoform HK1, are diminished, possibly due to neurodegeneration (Fig. S1G). Similar findings are reported by the Accelerating Medicines Partnership—Alzheimer’s Disease (AMP-AD) consortium, which comprises the currently largest collaborative post-mortem brain RNA-sequencing project. When the data is segregated by sex, females display an increase of HK2 in the dorsolateral prefrontal cortex and the parahippocampal gyrus (Fig. S1H). No brain region was significantly upregulated for HK2 expression in men (Fig. S1G. https://agora.ampadportal.org; accessed November, 2023).

Remarkably, co-expression network analysis provided by Agora, reveals that HK2 is up-regulated in the brain of AD patients in concert with several microglial genes (77% of the network, Fig. S1J) associated with neurodegeneration, including C1q, C3^22^, spp1^23^ and TLR2^24^, among others (Fig. 1J). Gene ontology analysis show that the main biological process associated with this “HK2/hAD network” involves immune responses and regulation of metabolic process (Fig. 1K) suggesting a key role of human HK2 as a hub between metabolism and microglial function. These results suggest that the microglial upregulation of HK2 observed in the 5xFAD mice is conserved in patients with AD and represents a potential therapeutic target. The sex difference observed in HK2 upregulation, is consistent with the hypothesis that microglial metabolism plays a central role in the sexual dimorphism of AD ^25,26^ and highlight the need to include both sexes in the assessment of potential disease modifying agents.

### Gene dosage of microglial HK2 selectively modulates the immune response and disease progression

To directly dissect the role of HK2 in the control of microglial immune response during AD progression, we generated mice that have a conditional deletion of one or two copies of HK2 gene in microglial cells by crossing 5xFAD;CX3CR1-Cre^ERT2^ mice with mice harboring a floxed HK2 allele^27^.

We induced the microglial deletion of one or both HK2 alleles (termed 5xFAD;HK2^Fl/wt^ or 5xFAD;HK2^Fl/Fl^, respectively) by tamoxifen (TAM) i.p. injections at 2 months old (Fig. 2A), a period at which amyloid deposition and microgliosis begin^28^. Mice harboring the Cx3cr1-Cre^ERT2^ cassette, but lacking the LoxP sites in HK2, were treated with TAM and used as controls (5xFAD;HK2^wt/wt^). Three months later (5 months of age), we evaluated AD pathogenesis (Fig. 2A).

**Fig. 2.**
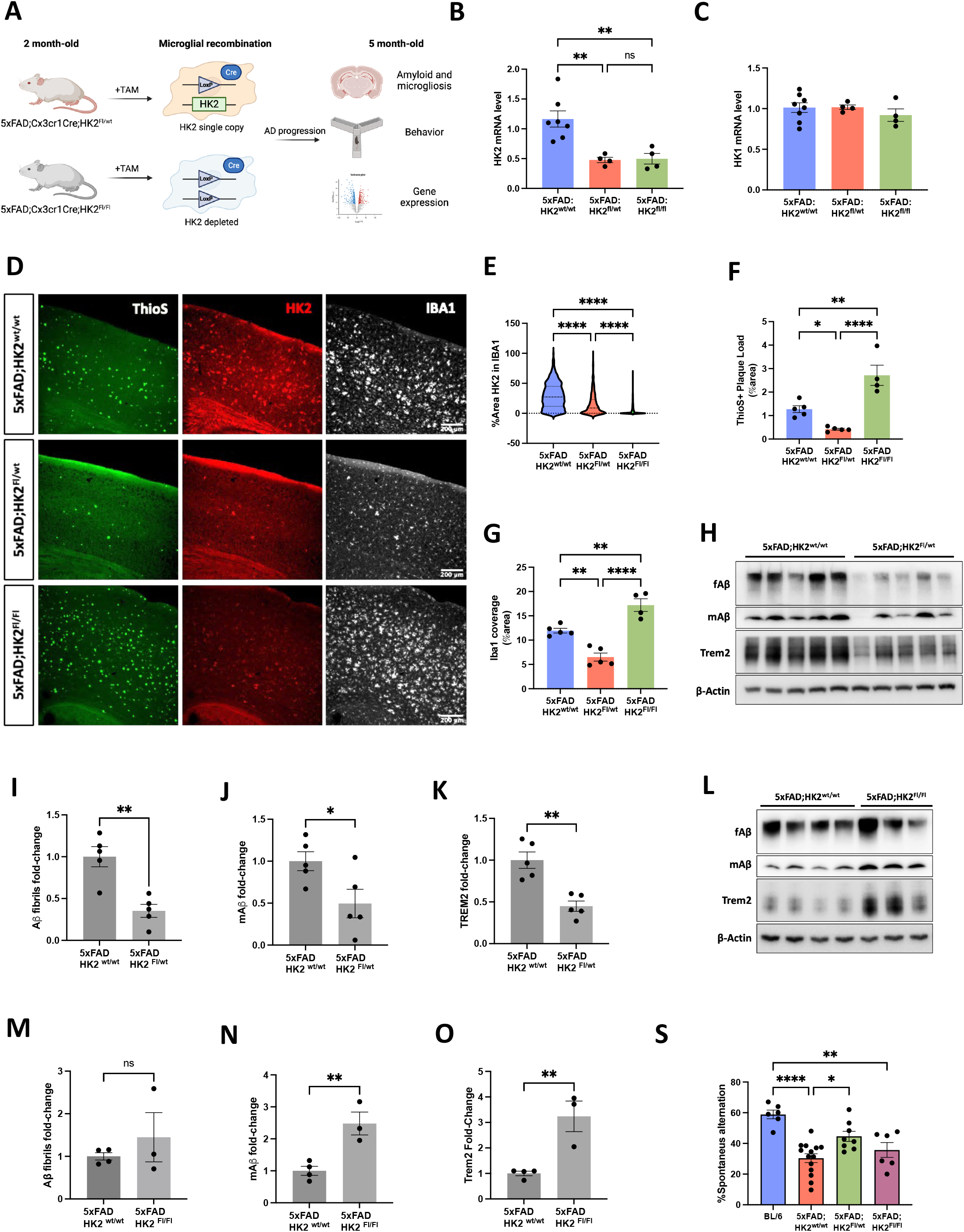
Gene dosage of microglial HK2 selectively modulates disease progression in the 5xFAD mice. **(A)** Mice that have a conditional deletion of one or two copies of HK2 gene in microglial cells were generated by crossing 5xFAD;CX3CR1-Cre^ERT2^ mice with mice harboring floxed HK2 alleles. At 2 months, five i.p. injections of TAM were administrated to induce recombination. Mice were assessed at 5 months old. **(B and C)** qPCR analysis of HK2 and HK1 expression in the cortex of 5xFAD mice harboring two (HK2^wt/wt^), one (HK2^Fl/wt^) or no copies (HK2^Fl/Fl^) of HK2 gene. **p< 0.01, *p< 0.05 (n=4-7 per group, One-way ANOVA followed by Tukey’s post-hoc test). **(D-G)** Immunofluorescent analysis of HK2 (red), microglia (Iba1, white) and Aβ (ThioS, green) in the cortical region of control and HK2 deficient mice. **E.** Quantification of the percentage of microglial area containing HK2 immunoreactivity shown in D, n=12 slices from 4 to 5 mice. **F and G.** Quantification of cortical area fraction of ThioS (plaque load) and microglia (Iba1^+^ cells). *p< 0.05, **p< 0.01, ****p< 0.0001 (n=4-5 per group, One-way ANOVA followed by Tukey’s post-hoc test). **(H)** Representative immunoblots of fibrillar and monomeric Aβ detected by MOAB2 antibody and Trem2 from soluble fraction of cortical lysates of 5xFAD;HK2^wt/wt^ and 5xFAD;HK2^Fl/wt^ mice treated with TAM. **(I-K)** Immunoreactive bands were quantified, normalized to β-actin and expressed as the fold change respect to 5xFAD;HK2^wt/wt^ mice. *p< 0.05 and **p< 0.01 (n=5, Unpaired t-test). **(L)** Representative immunoblots of fibrillar and monomeric Aβ detected by MOAB2 antibody and Trem2 from soluble fraction of cortical lysates of 5xFAD;HK2^wt/wt^ and 5xFAD;HK2^Fl/Fl^ mice treated with TAM. **(M-O)** Immunoreactive bands were quantified, normalized to β-actin and expressed as the fold change respect to 5xFAD;HK2^wt/wt^ mice. *p< 0.05 and **p< 0.01 (n=3-4 per group, Unpaired t-test). **(S)** Percentage of spontaneous alternation behavior in a Y-maze as an evaluation of spatial working memory. (n=6-14 per group, One-way ANOVA followed by Tukey’s post-hoc test)

HK2 deletion was confirmed by IHC (Fig. 2D and E) and qPCR with no changes observed in the ubiquitous HK1 (Fig 2B and C).

We evaluated the effect of microglial HK2 gene dosage on the cortical levels of Aβ. ThioS staining revealed that microglia haploinsufficient in HK2 induced a robust reduction in the dense core plaque burden in comparison with its HK2 sufficient littermates (Fig. 2D and F). Similarly, we observed a significant reduction in the microglial coverage area (Iba1^+^ cells), as a proxy for microglial density (Fig 2D and G). Remarkably, the complete deletion of HK2 resulted in an opposite phenotype from that observed in mice haploinsufficient for HK2, as the HK2 null mice were characterized by increased number of plaques and plaque associated microglia (Fig 2D, F and G). Consistently, western blot analysis of soluble fractions of the cortex of our transgenic mice mirrors the diverging effect of HK2 copy number in the levels of species of Aβ detected with MOAB2 antibody, as well as the microglial marker, Trem2 (Fig. 2H-O).

To evaluate if alteration in the expression of HK2 affects cognition, we evaluated working memory of the 5xFAD mice, as assessed by their spontaneous alternation in a Y-maze^29^. The partial loss of HK2 induced a significant improvement in the cognitive performance of the mice (Fig. 2S). Once again, the loss of both HK2 alleles had little effect on cognition, as the spontaneous alternation rate was similar to the control 5xFAD;HK2^wt/wt^ mice (Fig. 2S). No obvious sex differences were observed upon genetic inactivation of the HK2 gene.

To ascertain the cellular and molecular pathways affected by HK2 deficiency during AD progression, we performed transcriptional analysis of the cortical samples with NanoString at 5 months of age. We used a glia panel, that evaluates the expression of 770 genes, allowing us to perform gene set enrichment analysis (GSEA) to quantify cell populations defined by their corresponding marker gene set in each individual sample. The reduction in gene dosage of HK2 resulted in dramatically different outcomes, with HK2 haploinsufficiency shifting the expression of microglial genes towards homeostatic phenotypes (Fig. 3A). Conversely, the loss of both HK2 alleles had no effect on the transcriptional landscape of the 5xFAD microglia. With a less conservative analysis, we only observed changes in 3 genes that point to a detrimental effect, as the mitochondrial Arginase type 2 (Arg2) was decreased. α-synuclein and KLK6, a protease that participates in the cleavage of APP and were up-regulated. (Fig. 3B).

**Fig. 3.**
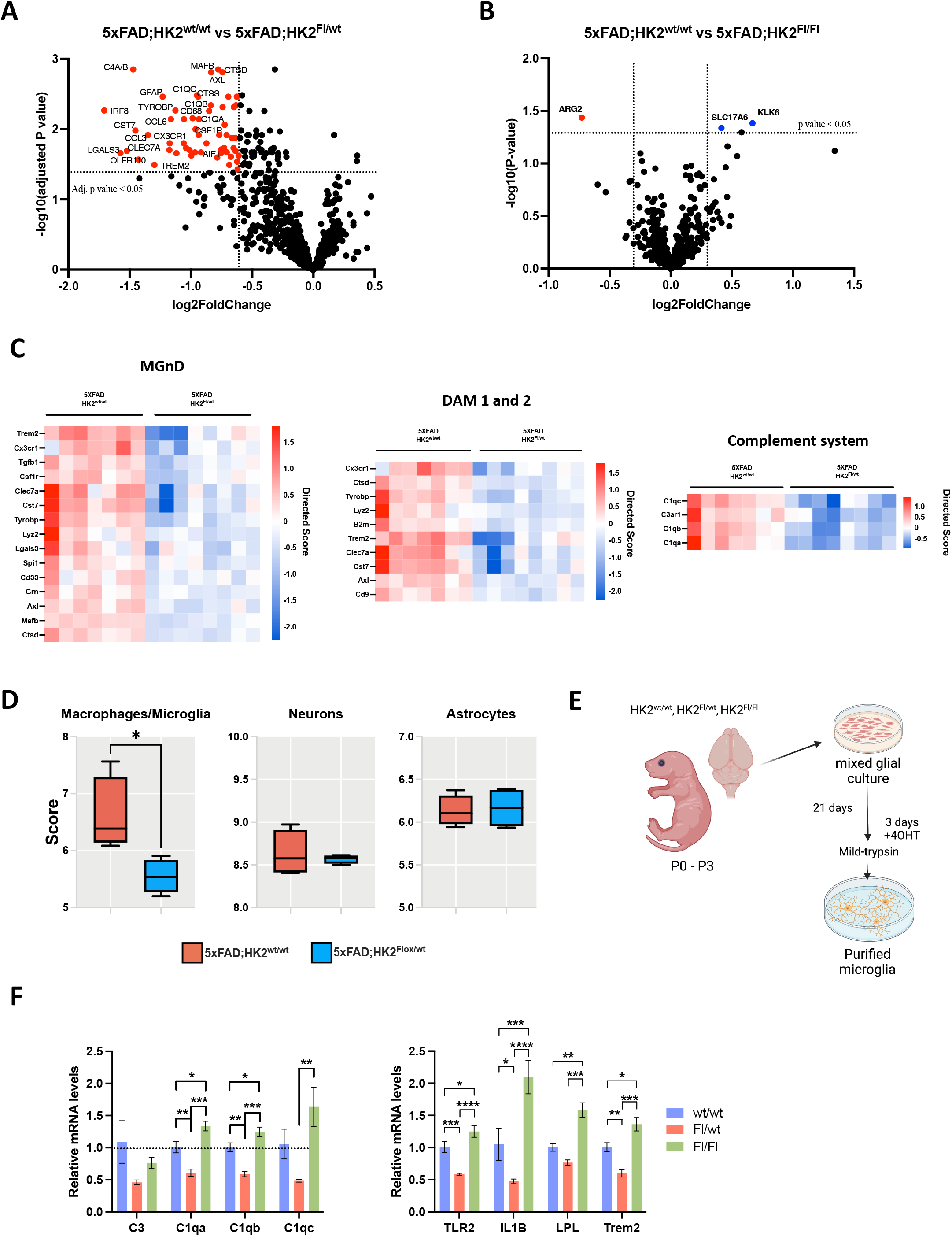
Gene dosage of microglial HK2 selectively modulates microglial gene expression in the 5xFAD mice. **(A and B)** Effect of HK2 gene dosage on gene expression respect to 5xFAD;HK2^wt/wt^ control mice. Differences for 770 genes in Nanostring glia panel are expressed as volcano plot. Adjusted P values were obtained from Rosalind on ramp software. Significance set at adj-pL<L0.05 and fold change >1.5, < -1.5 (n=7-8). **(C)** Heatmaps depicting the directed significance scores for genes annotated as markers of neurodegenerative phenotype (MGnD), DAM signature and complement system. The score magnitude of these gene sets was reduced only in the HK2 haploinsufficient AD mice. **(D)** Cell abundance scores reveal a microglial specific effect of HK2 partial loss. Cell type scores are based on the NanoString Cell Type Profiling Module *p< 0.05 (n=5, Unpaired t-test). **(E)** Schematic of the experimental procedure for the analysis of primary cultures of microglia. 4-OHT (4-hydroxytamoxifen). **(F)** qPCR analysis of selected microglial genes in primary cultures of HK2-deficient microglia. *p< 0.05, **p< 0.01, ***p< 0.001, ****p< 0.0001 (n=4-5 per group, One-way ANOVA followed by Tukey’s post-hoc test).

The effect of HK2 haploinsufficiency impacted the enrichment of gene sets associated with microglial activation and the transition to neurodegenerative states. GSEA of 5xFAD;HK2^Fl/wt^ mice showed a reduction in the MGnD/DAM signature genes as well as the complement system genes (Fig. 3C). Importantly, cell type profiling showed that the only cell type affected by HK2 partial loss of function was microglia, confirming the cellular specificity of our mice model and reinforcing the idea that the antagonism of HK2 attenuates the microglial response during AD progression without notable effects in other cells in the brain (Fig. 3D). This reduction in the enrichment of microglia is consistent with the reduced microglial abundance as demonstrated by IHC (Fig. 2G).

To directly evaluate the transcriptional effect of HK2 dosage on the microglial gene expression, we performed incubations of primary cultures of microglia derived of our transgenic mice with the active metabolite of TAM (4-OHT) (Fig. 3E). In this experimental setting we corroborated that a set of genes associated with immune activation and inflammation, (including LPL, IL1β, Trem2, TLR2, and genes of the complement system) exhibited reduced expression in the HK2 haploinsufficient microglia (Fig. 3F). Remarkably, the same set of genes displayed an increase, respect to controls, in the double KO HK2 microglial cells (Fig. 3F).

Taken together, our results suggest the HK2 expression level governs microglial phenotypes regulating key disease related responses such as microglial gene expression affecting both deposited and monomeric amyloid. This analysis is remarkable as the effects of HK2 gene dose are in opposite directions.

### Pharmacologic inhibitor of HK2, Lonidamine, modulates the microglial response and amyloid pathology in a sex dependent manner

To assess the therapeutic potential of HK2 inhibitors, we evaluated the acute effect of its pharmacological inhibition in the 5xFAD mice. Several drugs have been developed as HK2 specific inhibitors^30^. The best characterized is Lonidamine (LND) an orally administered small molecule that inhibits glycolysis by the preferential inactivation of HK2^31^. LND has been used in a number of clinical trials in cancer in the US and is currently approved in Europe for this indication^30^. LND is a selective blocker of glycolysis in tumor cells with minimal effects on normal cells and an acceptable side effect profile^32^.

We treated 5xFAD mice at different stages of disease for 7 consecutive days with a daily i.p. injection of 50 mg/kg of LND. Control mice (BL/6 and 5xFAD littermates) were injected with vehicle. The treatment did not affect the body weight of mice during the treatment period (Fig. S2A).

In males at 5 months of age, LND treatment induces a significant reduction in the cortical levels of fAβ, as revealed by ThioS staining (Fig. 4A and B). Similarly, we observed a significant reduction in the microglial coverage area (IBA1+ cells), reflecting a decrease in microgliosis (Fig 4C). However, the remaining plaques observed in the cortex, retain abundant plaque associated microglia and sustained barrier integrity as the microglial coverage of Aβ-plaques was not different between groups (Fig. 4D). The preservation of the barrier formation was functionally validated by evaluation of the area occupied by swollen Lamp1^+^ dystrophic neurites, which was reduced in the male LND group, suggesting an improvement in neuronal integrity, despite the overall decreased microgliosis (Fig. 4A and E).

**Fig 4.**
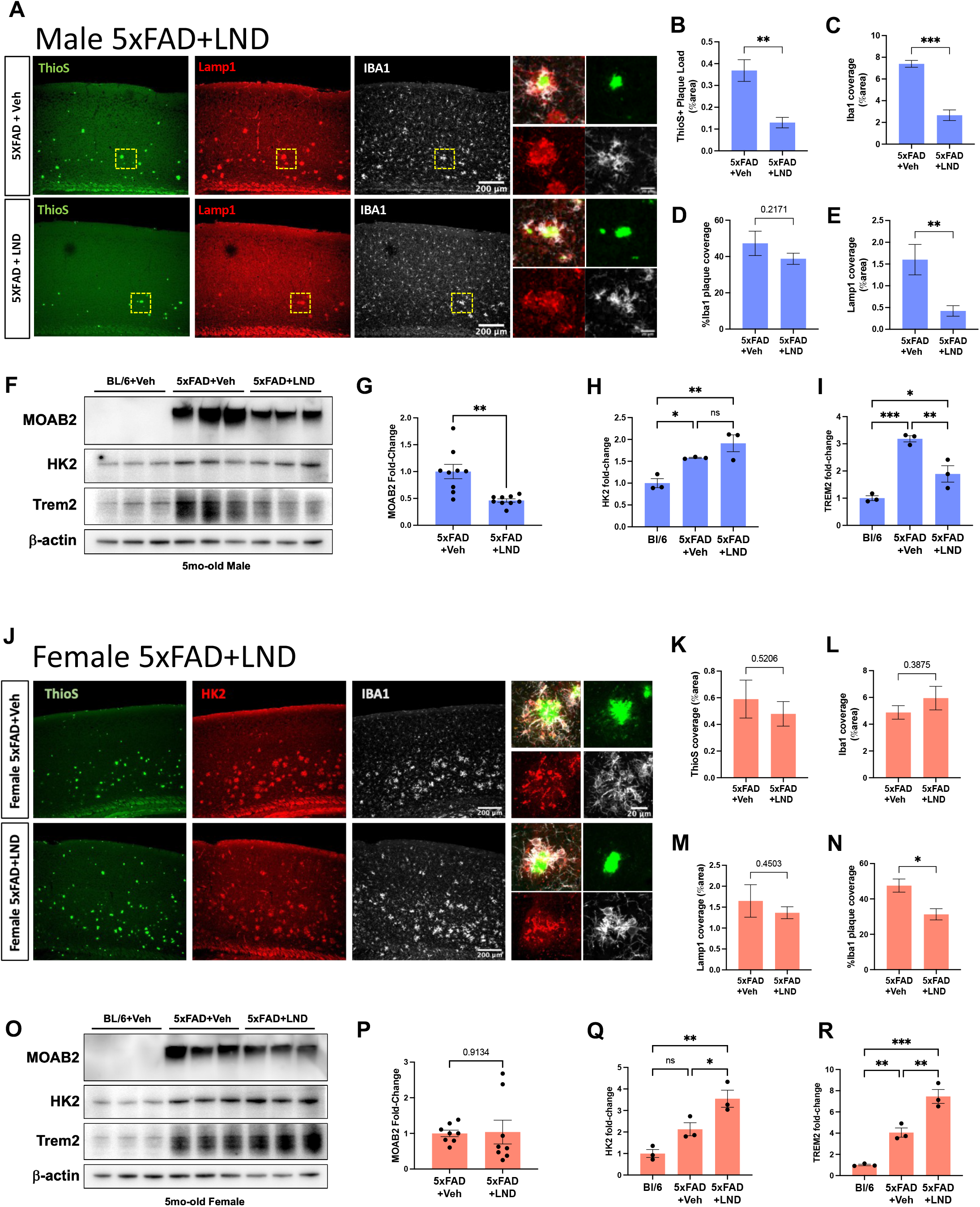
HK2 inhibition with Lonidamine, modulates amyloid pathology in a sex dependent manner. **(A)** Immunohistochemical analysis of the cortical region of 5-mo-old 5xFAD male mice treated with LND or vehicle. Amyloid plaques were stained with ThioS (green), dystrophic neurites with Lamp1 (red) and microglia with Iba1 (white). Dashed square area was magnified to better visualize microglial plaque coverage and Lamp1 area (right panels). **(B-E)** Quantification of cortical area fraction of ThioS (B), Iba1 (C) and Lamp1 (D). (E) Microglial plaque coverage was evaluated by quantify the average percentage area of Iba1 in individuals ThioS plaques across the whole cortex. (n=8-10 per group, Unpaired t-test) **(F-I)** Western blot and quantification of proteins from soluble fraction of cortical lysates of BL/6 and 5xFAD male mice treated with vehicle or LND. (F) Representative immunoblots of fibrillar Aβ detected with MOAB-2 antibody, microglial proteins HK2, and Trem2. (G-H) Immunoreactive bands were quantified, normalized to β-actin and expressed as the percentage of 5xFAD +veh mice in the case of MOAB-2. **p< 0.01 (n=9, Unpaired t-test). For the other proteins the values are expressed as the percentage of BL/6 +veh mice. (*p< 0.05, **p< 0.01, ***p< 0.001. n=3, One-way ANOVA, Tukey posttest). **(J)** Immunohistochemical analysis of the cortical region of 5-mo-old 5xFAD female mice treated with LND or vehicle. Amyloid plaques were stained with ThioS (green), dystrophic neurites with Lamp1 (red) and microglia with Iba1 (white). The dashed square area was magnified to better visualize microglial plaque coverage and Lamp1 area (right panels). **(K-N)** Quantification of cortical area fraction of ThioS (K), Iba1 (L) and Lamp1 (M). (N) Microglial plaque coverage was evaluated by quantify the average percentage area of Iba1 in individuals ThioS plaques across the whole cortex. (n=8-10 per group, Unpaired t-test) **(O-R)** Western blot and quantification of proteins from soluble fraction of cortical lysates of BL/6 and 5xFAD male mice treated with vehicle or LND. (O) Representative immunoblots of fibrillar Aβ detected with MOAB-2 antibody, microglial proteins HK2, and Trem2. (P-R) Immunoreactive bands were quantified, normalized to β-actin and expressed as the percentage of 5xFAD +veh mice in the case of MOAB-2. (n=8, Unpaired t-test). For the other proteins the values are expressed as the percentage of BL/6 +veh mice. (*p< 0.05, **p< 0.01, ***p< 0.001. n=3, One-way ANOVA, Tukey posttest).

Western blot analysis of the soluble fraction of the cortical lysate, corroborate the effect of HK2 inhibition on amyloidosis, as fibrillar Aβ42, was dramatically reduced in the male 5-mo-old 5xFAD mice treated with the HK2 inhibitor (Fig. 4F and G) and almost undetectable at 3-mo (Fig. S2B and C). We also observed a significant reduction in the levels of Trem2, suggesting attenuation of microglial activation (Fig. 4F and G). However, the 7 day treatment failed to reduce the cortical amyloid burden or Trem2 levels at 8-mo (Fig. S2F-I) which is consistent with the idea that microglial intervention is more effective during the early stages of disease^33,34^.

These findings are similar to those recently reported by Leng *et al.*, however that study only included 5XFAD males evaluated at 6 months^14^.

Unexpectedly, the same treatment, failed to reduce amyloid, microgliosis and neuronal dystrophy in the female cortex (Fig. 4J-N). Western blot analysis of the cortical lysates with the anti-Aβ antibody MOAB2 showed no changes in amyloid burden with the drug treatment but resulted in a significant increase in the levels of HK2 and Trem2, suggesting an increase in the activation state of microglia. This trend was also observed at 3-mo (Fig. S2B-E) suggesting that the observed differences between sex were not attributable to an excessive amyloid burden in females, as at 3-mo the cortical amyloid load is lower that the observed in 5mo-old male cortex^35^. Analogous to that observed in males, the acute treatment with LND failed to induce changes in amyloid levels in the female cortex at 8-mo (Fig. S2F-I).

To ascertain the cellular and molecular pathways affected by HK2 inhibition, we performed transcriptional analysis of the cortical samples with NanoString. Our bulk RNA analysis confirms the sex-biased phenotype of microglia induced by LND, as females exhibit increased levels of several microglial genes associated with induction of an immune response, including Trem2, Tyrobp, Spp1, CD68 and complement-related genes compared to males at 5 mo of age (Fig 5A). Complementary qPCR analysis corroborates these opposite changes for selected DAM markers (Fig. 5B). GSEA showed that the major differences in the biological processes between sex are related to their activation status, with females showing a higher expression of MGnD/DAM signature genes as well as the one associated to the primed microglia and the complement system (Fig. 5C). This observation clearly suggests that LND treatment is inducing sex specific effects in the microglial activation of 5xFAD mice. This was further supported by cell type profiling, that like our above mentioned IHC analysis (Fig. 4C and L), show a reduced microglial abundancy in males compared to females with no detectable changes in other cell types. These data also confirm the cellular specificity of HK2 and its targeting with LND (Fig. 5D).

**Fig 5.**
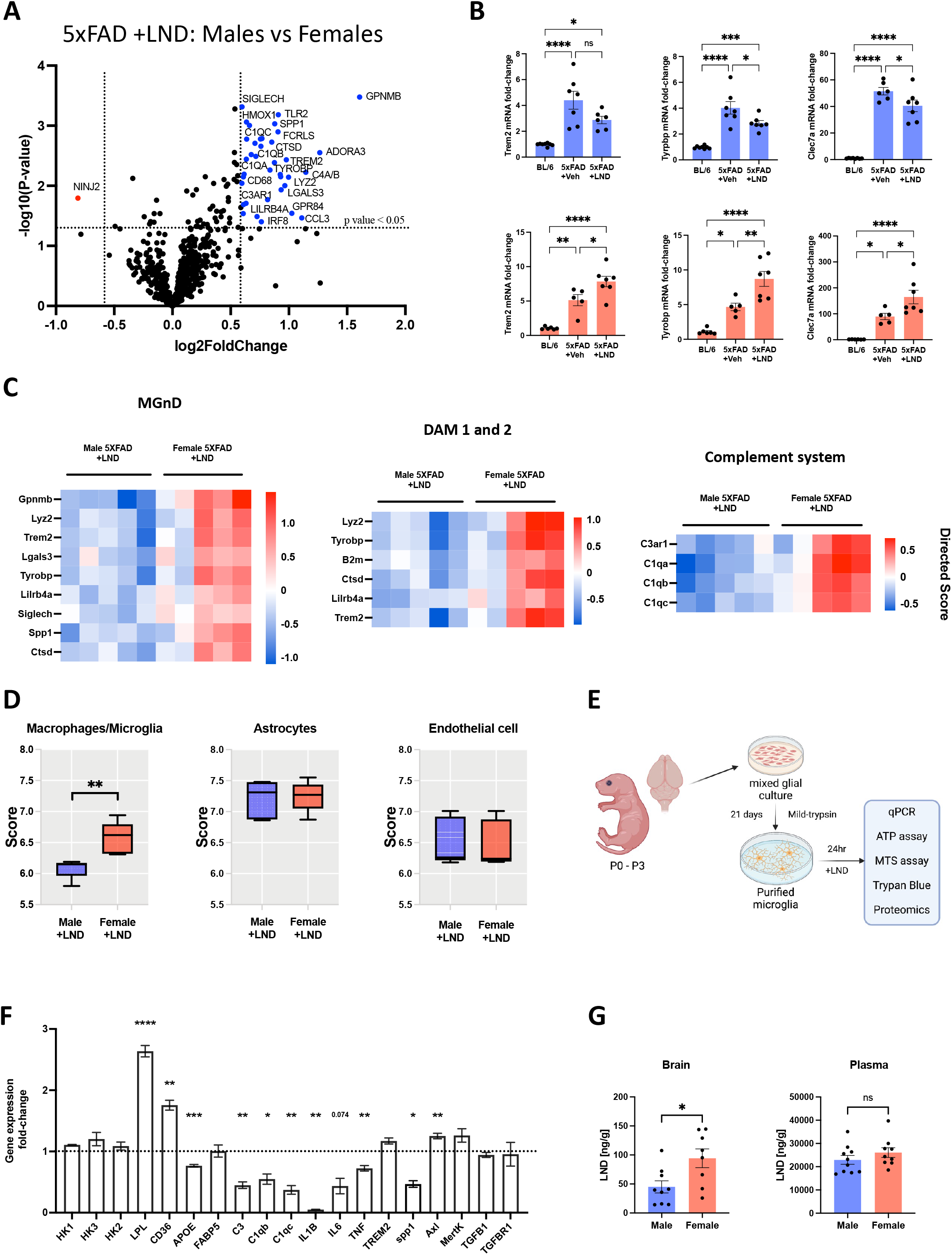
LND regulates the neuroinflammatory activation of microglia in a sex dependent manner. **(A)** Volcano plot illustrating statistical significance of sex-related differences in gene expression after treatment with LND in 5xFAD mice. Significance set at pL<L0.05 and fold change < 1.5 (n=5 per group). **(B)** qPCR analysis of selected microglial genes in the cortex of male (upper blue graphs) and female (lower, salmon graphs) 5xFAD mice treated with LND. *p< 0.05 and, **p< 0.01, ***p< 0.001, p<0.0001 (n=5-8 per group, One-way ANOVA, Tukey posttest). **(C)** Heatmap depicting the global significance scores for nanostring annotated pathways. GSEA revealed multiple immune associated pathways upregulated in the female cortex of 5-month-old mice treated with LND when compared to male treated mice. **(D)** Cell type abundance scores are based on the NanoString Cell Type Profiling Module. *p< 0.05 (n=5 per group Unpaired t-test). **(E)** Schematic of the experimental procedure for the analysis of primary cultures of microglia treated with LND. **(F)** qPCR analysis of selected microglial genes in primary cultures of microglia treated with LND. *p< 0.05 and, **p< 0.01, ***p< 0.001 (n=3 per group, Unpaired t-test). Dashed line represents gene expression in vehicle treated cultures. **(G)** Lonidamine brain and plasma levels evaluated by HPLC after 4 hrs. of a single i.p. injection. *p< 0.05 (n=8 and 9 per group Unpaired t-test).

This bulk analysis of gene expression could be affected by the altered number of microglia between sexes. To further evaluate the direct effect of LND on the microglial gene expression, energy production and viability we performed incubations with LND in primary cultures of microglia for 24 hrs (Fig. 5E). In this experimental setting, the sex effect was completely lost as we observed a significant and consistent reduction in the mRNA expression of inflammatory cytokines (IL1β, IL6 and TNF), complement proteins, as well as the DAM markers spp1 and APOE (Fig. 5F). Additionally, the expression of HK1 as well as homeostatic markers like TGFB1 and TGFBR1 were not affected (Fig. 5F), which is a profile comparable to the observed only in male treated *in vivo*.

Different factors have been proposed to explain the clinical sex differences observed in the incidence of AD, response to treatments and occurrence of side-effects^36^. Among these factors, sex differences in drug pharmacokinetics and brain exposure have been proposed to explain the two fold incidence of adverse drug reactions in women compared to men^37^. We compared the LND brain levels in both sexes, to determine potential differences in biodistribution that could explain the opposite microglial phenotypes observed *in vivo*. HPLC-MS/MS analysis found that in females the brain concentrations of LND were the double (94.29 ± 16.15 ng/g) that in males (45 ± 10.47 ng/g), despite comparable blood concentrations (Fig. 5G). This could also explain the lack of divergent effects observed *in vitro*. However, we cannot discount the effect of additional factors (i.e., hormones) in the induction of opposite phenotypes *in vivo,* as these too were absent in our *in vitro* studies. Currently, we do not know if this differential brain concentration is due to differences in BBB integrity, transport, or brain metabolism of LND. However, this finding suggests that the sex biased effect of LND on microglial phenotype may be dependent on the levels of HK2 activity in brain. Additional experiments are necessary to determine the basis of differential LND distribution in the brain and if an analogous situation is observed in humans.

### A protective phenotype of microglia induced by HK2 antagonism, is not associated with increased ATP production nor increased mitochondrial binding of HK2

Multiple mechanisms have been proposed to explain the role of HK2 in the modulation of inflammatory responses of microglia/macrophages, including its metabolic actions and newly recognized alternative mechanism arising from its regulated association with mitochondria.

It has recently been argued that microglial HK2 antagonism dramatically boosted ATP production exclusively within microglia through a compensatory shift to LPL-dependent lipid metabolism, increasing the phagocytosis of Aβ in murine models of AD^14^. However, a second independent study by Hu et al., arrived at quite different conclusions as they found that microglial HK2 depletion, resulted in reduced levels of cellular ATP and mitochondrial dysfunction, a setting which would not support increased mitochondrial β-oxidation of lipids^15^.

Because our phenotypic characterization showed that the beneficial effects in the 5xFAD mouse was observed upon partial antagonism of HK2, we evaluated ATP production in microglia haploinsufficient for HK2 or microglia treated with the inhibitor LND. In both conditions the HK activity was significantly reduced (Fig. 6A and B), however the *in vitro* measurements of ATP production show disparate results, with no changes in ATP levels observed in primary cultures haploinsufficient for HK2 (Fig. 6A). In LND-treated cells, we observed a dose-response decrease in the levels of ATP that was significant at 200 μM of LND (Fig 6B). Cellular metabolism was evaluated by MTT assay, which is based in the activity of mitochondrial NAD(P)H-dependent dehydrogenase enzymes in metabolically active cells^38^. No changes were observed with HK2 partial loss or between 50 and 200 μM of LND (Fig. 6A and B) suggesting that cells are viable and mitochondrial function was not affected in these conditions. Higher LND concentrations (>500 μM) elicited cellular toxicity which also reflects a massive drop in ATP levels (Fig. 6B). We conclude that the beneficial effects of HK2 partial antagonism are not dependent on elevated ATP production.

**Fig. 6.**
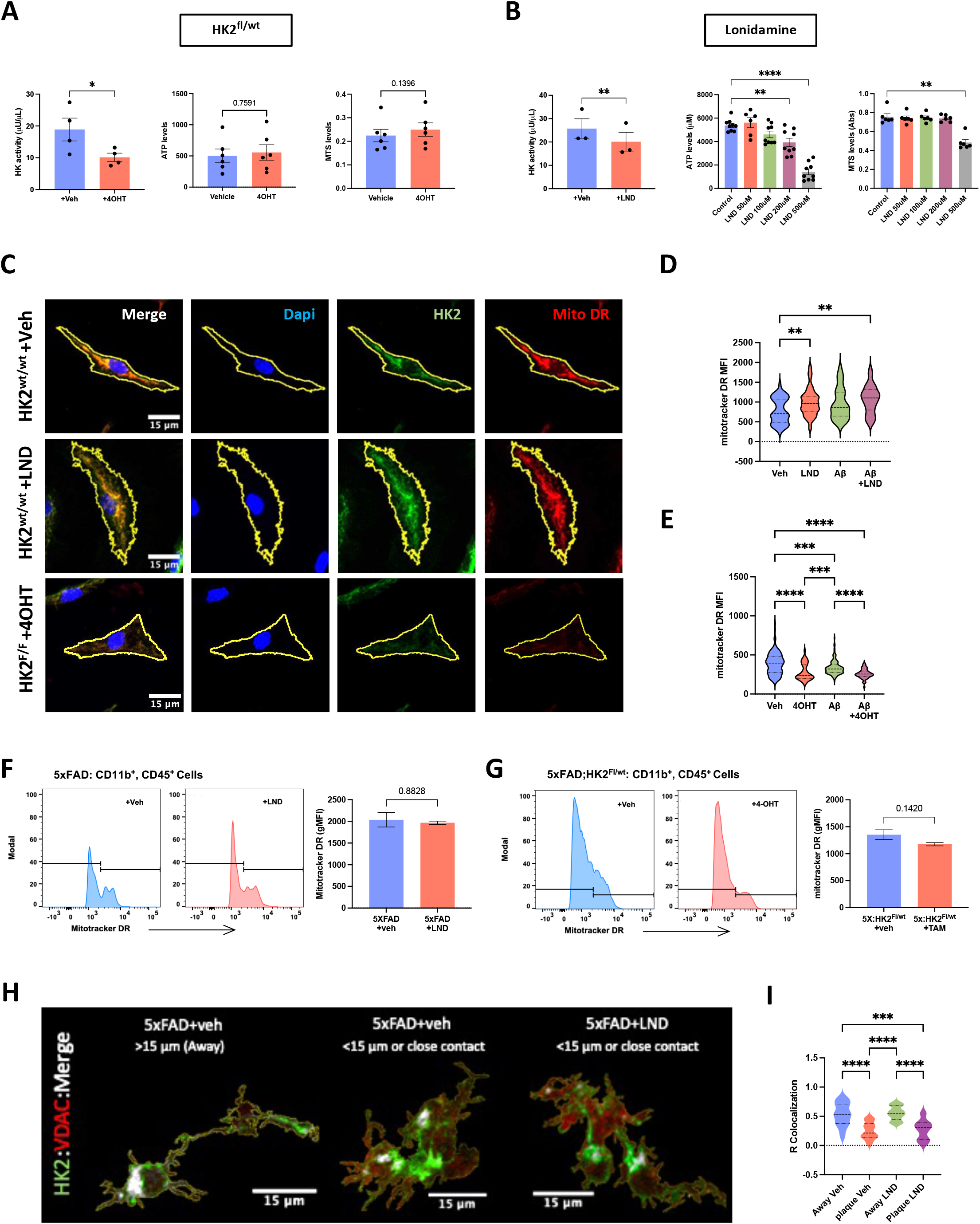
HK2 inhibition reduces ATP production and its gene dosage induce divergent effects on mitochondrial function. **(A)** *In vitro* evaluation of HK activity, ATP production and cellular metabolism in primary cultures of microglia haploinsufficient for HK2. (*p< 0.05, n=4-6 per group, Unpaired t-test) cultures **(B)** *In vitro* evaluation of HK activity, ATP production and cellular metabolism in primary cultures of microglia treated with different concentrations of HK2 inhibitor LND. (**p< 0.01, n=3 per group, Unpaired t-test and **p< 0.01, ****p<0.0001, n=9 per group, One-way ANOVA, tukey posttest, respectively) **(D and E)** Quantification of the mean fluorescence intensity (MFI) of mitotracker DR in primary microglia described in C. **p< 0.01, ***p<0.001, ****p<0.0001, n=9 per group, One-way ANOVA, tukey posttest, respectively) **(F)** Flow cytometry analysis of mitochondrial potential in CD11b^+^, CD45^+^ cells treated with LND. Left, representative histogram of the fluorescence intensity of mitotracker deep red (DR) in microglia from 5xFAD mice treated with vehicle or LND. Right, quantification of the geometric mean fluorescence intensity (gMFI) of mitotracker DR. (n=3 per group, Unpaired t-test) **(G)** Flow cytometry analysis of mitochondrial potential in CD11b^+^, CD45^+^ cells of 5xFAD HK2 haploinsufficient mice as described in F. (n=3 per group, Unpaired t-test). **(H)** Colocalization analysis of HK2 and VDAC in the 5xFAD mice treated with LND. Representative confocal microscopy images showing HK2 (green) VDAC (red) and it colocalization (white) in resting and activated microglia from mice treated with vehicle or LND. Pearson quantification shown that activated microglia display decreased levels of mitochondrial association and LND treatment fails to recover this effect. (***p< 0.001, ****p<0.0001, n=4, >50 cells per group, One-way ANOVA, tukey posttest)

We further explored the effect of HK2 antagonism on microglial mitochondria, by evaluating its membrane potential, which reflects the process of electron transport and oxidative phosphorylation, the driving force behind ATP production. In primary cultures of microglia derived from HK2 KO mice, staining with Mitotracker Deep Red revealed a significant decrease in its mitochondrial potential after HK2 deletion (Fig. 6C and E). However, LND treatment of primary microglia, did not affect this parameter (Fig. 6C and D).

*In vivo*, we evaluated the microglial mitochondrial potential by flow cytometry of CD11b^+^,CD45^+^ cells derived from brains of our 5xFAD mice. In 5-mo-old mice, neither HK2 haploinsufficiency, nor LND treatment had significant effects on the number of activated microglia positive for Mitotracker DR (Histograms) or in its fluorescent intensity (Fig. 6F and G), implying that the partial antagonism of HK2 attenuates its catalytic activity without the mitochondrial side-effects observed with its total deletion. These results suggest that the differential effect of HK2 gene dosage on mitochondrial dysfunction, which is an important trigger of the inflammasome activation, could explain the opposite microglial response observed in AD, as consequence of HK2 copy number.

Finally, we studied the *in vivo* colocalization of HK2 and VDAC in the 5xFAD mice. In 5month-old mice we observed a significant decrease in HK2 mitochondrial association in plaque associated microglia compared to unaffected parenchymal microglia (Fig. 6H and I). As this phenomenon has been proposed as an important inflammatory trigger, we expected that LND treatment might revert the amyloid induced cytosolic translocation of HK2. However, LND failed to revert this pathological effect (Fig. 6H and I), arguing that the mechanisms by which HK2 partial antagonism regulates inflammation and disease progression are independent of its mitochondrial association and could be due to the inhibition of HK2 cytosolic actions.

### HK2 partial antagonism reduces the expression of elements of the inflammasome and the A**β**-induced nuclear-translocation of NFK**β**

Microglia are a multifaceted cell that participate in AD pathogenesis through many distinct mechanisms including phagocytosis and clearance of Aβ as well as the induction of neuroinflammation and the spreading of Aβ^39^. Primary cultures of microglia, treated with LND increased their uptake of Aβ, as revealed by WB and MOAB2 staining. (Fig. 7A and B) as previously reported (Leng). However, flow cytometry of microglia revealed that LND did not affect the percentage of phagocytic cells or levels of internalized Aβ, evaluated by Methoxy-X04 uptake (Fig. 7C). Additionally, we observed that our manipulations of HK2, failed to alter the amyloid burden in brain areas experiencing early accumulation, like the subiculum (Fig. S3A-D), or when the treatment is done in the symptomatic stage of the disease (8-mo-old, Fig. S2F-I). This suggests that the phagocytic capacity induced by HK2 partial inhibition is probably necessary but not sufficient to eliminate the pre-existing amyloid plaques.

**Fig. 7.**
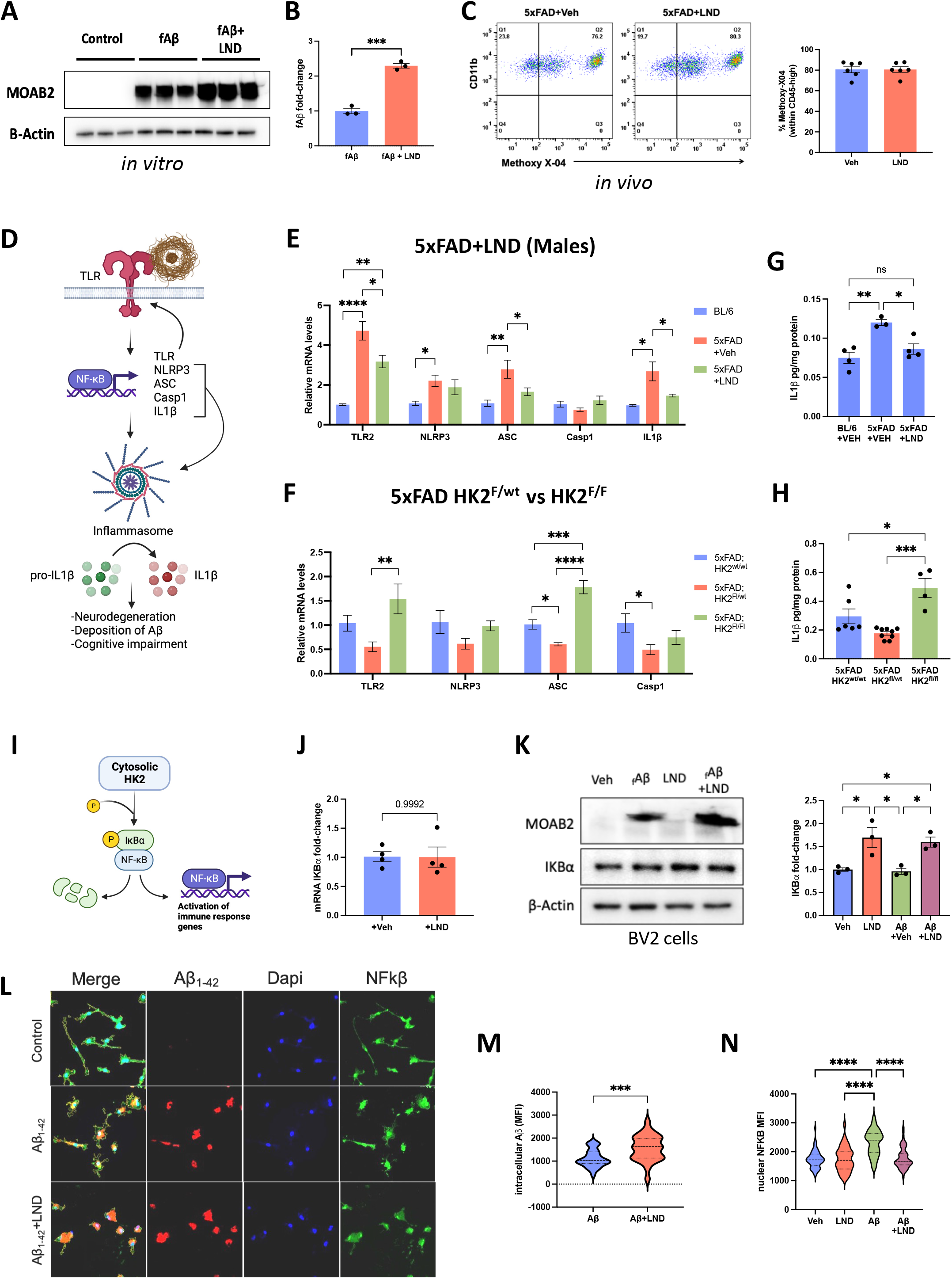
HK2 gene dosage induces opposite results in the induction of inflammation. **(A and B)** Western blot analysis of fibrillar Aβ (MOAB2) in primary cultures of microglia co-incubated with aggregated Aβ_1-42_ and LND for 24 hrs. (n=3, Unpaired t-test) **(C)** Flow cytometry analysis of methoxy X-04 uptake in CD11b^+^, CD45^+^ microglia from 5xFAD mice treated with vehicle or LND. No changes were observed in the percentage of phagocytic cells (n=6, Unpaired t-test). **(D)** Schematic of the inflammasome pathway and the production of IL-1β. **(E)** qPCR analysis of elements of the inflammasome in males Bl/6, 5xFAD and 5xFAD treated with LND. *p< 0.05, **p< 0.01, ****p< 0,0001 (n=4-6 per group, One-way ANOVA, tukey posttest). **(F)** qPCR analysis of elements of the inflammasome in 5xFAD mice with different gene dosage. *p< 0.05, **p< 0.01, ***p< 0.001, ****p< 0,0001 (n=4-5 per group, One-way ANOVA, tukey posttest). **(G and H)** ELISA analysis of IL-1β levels in LND treated males 5xFAD mice (G, *pL<L0.05, n=3-4, One-way ANOVA, tukey posttest) and in 5xFAD mice with different HK2 gene dosage. *pL<L0.05, ***p<0.001 (H, n=4-9, One-way ANOVA, tukey posttest) **(I)** Schematic of HK2 cytosolic regulation of NFKβ nuclear translocation mediated by its actions over IKBα protein degradation as described by Guo *et al*. (A) **(J)** qPCR analysis of IKBα mRNA expression in primary cultures of microglia treated with LND. (n=4, Unpaired t-test). **(K)** Western blot analysis of IKBα in primary cultures of microglia co-incubated with aggregated Aβ_1-42_ and LND for 24 hrs. *p<0.05 (n=3, One-way ANOVA, tukey posttest) **(L)** Representative confocal microscopy images showing NFKβ (green) nuclear translocation (dapi, blue) in primary cultures of microglia co-incubated with aggregated Aβ_1-42_ (red) and LND for 24 hrs. **(M and N)** Quantification of intracellular Aβ and nuclear NFKβ mean fluorescence intensity as shown in L. ***p<0.001, ****p< 0,0001 (n=3 cover slips from 3 independent cultures, Unpaired t-test and One-way ANOVA, tukey posttest, respectively)

It has been proposed that inflammation can impact amyloid formation by the seeding activities of some elements of the inflammasome, like ASC^40^. Additionally, the production of pro-inflammatory factors which act to remodel synapses by altering the shape, composition, and density of synapses that untimely contribute to the cognitive dysfunction of AD patients^41^ (Fig. 7D).

Accordingly, we evaluated the expression of the components of the inflammasome pathway described in Fig 7D. qPCR analysis has shown that at 5 months, most of the elements of the inflammasome are upregulated in the cortex of the 5xFAD mice (Fig. 7E and S3E). In accordance with our previous observations, the treatment with LND induced a sex-biased effect. In males, HK2 inhibition blocked the increase for TLR2, ASC and the inflammatory effector IL1β (Fig. 7E). Female mice display a polar opposite phenotype with increased expression of inflammasome components in response to HK2 pharmacological inhibition (Fig. S3E and F).

As reported above, the sex differences in gene expression were loss in our genetic approach, but we still observed a HK2 gene dosage effect. 5xFAD mice haploinsufficient for HK2 shown a reduction in the expression of inflammasome elements, meanwhile the double KO display an increase in the levels of TLR2 and ASC (Fig. 7F). Elisa analysis confirms the reduction of the protein levels of IL1β after LND treatment in males (Fig. 7G) and the divergent effect induced by HK2 gene dosage (Fig. 7H).

Recent reports have suggested that HKs can induce inflammation by a gain of non-metabolic functions associated with the induction of the inflammasome through the NF-Kβ pathway^16,37^. NF-Kβ is a master regulator of innate and adaptive immune functions that plays a critical role in the expression of various proinflammatory genes including inflammasome regulation^42^ (Fig. 7D). Moreover, NF-Kβ has been described as a downstream target of metabolic sensors and its transcriptional activity can be regulated by inhibitors of glycolysis in microglia challenged with LPS ^43^. We explored a novel HK2-dependent mechanism, recently described in cancerous cells^44,45^. This mechanism involves a cytosolic gain of function for HK2 in which the binding and phosphorylation of IKBα, a cytosolic repressor of NFKβ, results in IKBα degradation and subsequent nuclear translocation of NFKβ (Fig. 7I). Primary cultures of microglia were treated with aggregated Aβ and LND for 24 hrs to determine the levels of IKBα and nuclear translocation of NF-Kβ. qPCR analysis revealed that HK2 inhibition has no effect on IKBα transcription (Fig. 7J), however western blot analysis shown that LND treatment significantly increased its protein levels even in the presence of Aβ (Fig. 7K). As expected, Aβ treatment induced increased presence of NF-Kβ in the nuclei of challenged cells with respect to vehicle controls, however the co-incubation with LND prevent this translocation (Fig.7L and N). concomitant with an increase in the uptake of Aβ (Fig.7L and M). This suggest that HK2 role in the stabilization of IKBα and NF-Kβ signaling is conserved in microglia. These findings substantially expand our understanding in the crosstalk between metabolism and inflammation in the context of neurodegenerative diseases, particularly AD.

Together, these results, point to inflammation as a possible mechanism by which HK2 dosage results in opposing functional phenotypes of microglia, influencing AD progression. We argue that the partial antagonism of HK2 regulates microglial responses by affecting the induction of inflammation by decreasing the NF-Kβ mediated expression of elements of the inflammasome and IL1β, this process seems to be dependent on its catalytic activity and not dependent on the energetic status of the cell. Remarkably, the partial antagonism of HK2 does not affect the mitochondria, suggesting that the mitochondrial dependent inflammatory pathway was not affected. This is not the case for the complete deletion of HK2, in which the induced mitochondrial deficits not only are inconsistent with increased ATP production, but also explain a hyper-inflammatory phenotype that exacerbates AD progression.

## Discussion

Central to understanding the biological underpinning of the microglia actions in the AD brain is the recognition of the importance of disease-induced reprogramming of cellular metabolism. Immunometabolism governs the ability of microglial to mount an immune response and the nature of its effector functions ^6,8,46^. Microglia respond to an immune challenge (in this case, exposure to deposited amyloid) by switching from oxidative metabolism to glycolysis to support the acquisition of an disease-induced phenotypic states typified by generation of complex and sustained immune responses^47^.

In this study we tested the hypothesis that targeting key enzymes within metabolic pathways that are employed by immune stimulated microglia might provide new therapeutic options for AD. We focused our attention on HK2, because it is a member of a family of enzymes that perform the first and rate limiting step in glycolysis through the irreversible phosphorylation of glucose which can be further metabolized either by glycolysis to rapidly generate ATP as well as other metabolic intermediates necessary for various biosynthetic pathways^12^. Recent evidence has demonstrated that HKs can regulate inflammation through both metabolic and non-metabolic activities^48^.

During the preparation of this manuscript, two independent groups performed analogous studies but reported conflicting data with respect to the effect of HK2 complete deletion and inhibition on microglial ATP levels and microglial responses in murine models of stroke and AD respectively^14,15^. This discrepancy is relevant as Leng *et al*, proposed that the increased ATP levels, through a compensatory shift to lipid metabolism is instrumental for AD improvement after microglial HK2 inhibition. This proposal however, conflicts with previous studies made in microglia treated with HK2 inhibitors that describe decreased levels of ATP. Cheng *et al*, describe that 2-Deoxy-D-glucose (2-DG) and 3-BP induce an increase in the ADP/ATP ratio of BV2 microglia which was associated with decreased inflammation^16^. Vilalta and Brown also observed that 2-DG treatment resulted in a rapid depletion of ATP that precede microglial apoptosis^49^. Hu *et al*, demonstrated that the treatment with 2-DG, 3-BP or the induced deletion of microglial HK2 resulted in a dramatic reduction of ATP, leading to an *“energy-deficient state in microglia, with no compensation to glutamine or fatty acid oxidation”*^15^.

Our results align better with those reported by Hu and others as we observed that the antagonism of HK2 results in reduced levels of cellular ATP. Another substantial difference, reported *in vivo*, was the inflammatory consequence of HK2 complete deletion in neurodegenerative contexts. Leng reports decreased inflammation and disease improvement in the 5xFAD mice model, meanwhile Hu, *et al*. demonstrated that HK2 complete deletion, exacerbate disease presentation in a stroke mice model, mainly due to an over inflammatory microglial response. Once again, our results align better with Hu, *et al.* as we observed that the double KO of HK2 failed to improve cognition and reduce amyloid in the 5xFAD mice. This was associated with a microglial phenotype characterized by elevated levels of elements of the inflammasome, the complement and IL1β production.

Hu *et al*, proposed that the complete deletion of HK2 induce mitochondrial dysfunction as one of its non-metabolic functions is the occlusion of the mitochondrial pore by its binding with Vdac^15^. Similar consequences were also reported in HK2 depleted epithelial cells, with drastically reduced mitochondrial respiration indicating impairments of the mitochondrial electron transport chain^50^. We confirmed that the double KO of HK2 resulted in a significant reduction in the mitochondrial potential which is directly linked with a reduced capacity of mitochondria to produce ATP. These joined observations are inconsistent with the mechanism proposed by Leng and colleagues as mitochondria health is fundamental for the ATP production associated to β-oxidation of lipids. Additionally, in macrophages, the lack of HK2 mitochondrial binding is a strong inflammatory trigger as allow the release of mitochondrial content that activate the inflammasome and stimulate the production of inflammatory cytokines^17^. This is consistent with the observed expression of elements of the inflammasome and the increased production of IL-1β observed in our HK2 double KO mice.

We expanded these studies by characterizing the HK2 heterozygous mice in the 5xFAD context, as we reasoned that a single copy of HK2 can regulate its inflammatory activities without compromising its mitochondrial role. Indeed, the partial loss of HK2 induced an opposite microglial phenotype characterized by reduced inflammation and amyloid burden as well as improved spatial memory. Importantly, neither the partial loss of HK2 nor LND treatment results in mitochondrial alterations, suggesting that these manipulations of HK2, specifically targets its catalytic activity without significant effects in its non-metabolic function related to mitochondrial integrity. This separation of roles is impossible for the complete loss of HK2 as its mitochondrial function depends on its molecular association with the mitochondrial pore in the outer membrane.

In our hands, HK2 partial antagonism was not inducing a boosted production of ATP nor restoring homeostatic association with Vdac. We explored a third mechanism, recently described in cancerous cells^44,45^. This mechanism involves a cytosolic gain of function for HK2 in which the binding and phosphorylation of IKBα, a cytosolic repressor of NFKβ, results in IKBα degradation and subsequent nuclear translocation of NFKβ. Consistently, HK2 partial antagonism resulted in elevated protein levels of IKBα that blocked the nuclear translocation of NFKβ. This suggests that this novel non-metabolic role of HK2 is conserved in microglia, highlighting its multifaceted role in the regulation of inflammation, particularly during AD progression.

Logically, the final phenotypic output of microglia is going to be determined by the balance between these different mechanisms regulated by HK2. The complete loss of HK2 represent and extreme in which mitochondrial dysfunction incline the balance towards inflammation and its partial deficiency toward homeostasis due to reduced NFKβ signaling. This phenomenon has precedent, as the group of Li Gan reported similar observations induced by Trem2 gene dosage on microglial injury response and tauopathy^51^. PLCG2, another important AD risk factor can regulate divergent microglial functions via Trem2 or TLRs association^52^, reinforcing the idea that many molecular players can drive microglial phenotypes depending not only on it enzymatic activity, but also in attention to its cellular localization and interacting partners.

This phenomenon represents both an opportunity and a challenge, as the pharmacological targeting of HK2 can also render opposite results as it was observed between males and females. We hypothesize that the difference is due to a higher drug concentration in the brain of females as revealed by HPLC, inducing a phenotype that resemble the observed in the double KO. Because women are nearly twice as likely as men to develop AD, include both gender in the assessment of new targets for AD therapeutics if fundamental as gender is one of the most important phenotypic variables observed in many failed clinical trials^53^. Determine the exact causes of this sex biased response will be instrumental to better understand how the molecular context determine HK2 activities in females and stablish safe metabolic interventions for all patients of AD.

## Material and methods

### Resource availability

#### Lead contact

Further information and requests for resources and reagents should be directed to and will be fulfilled by the Lead Contact, Gary E. Landreth

#### Materials availability

This study did not generate new unique reagents.

### Experimental Model and Subject Details

#### Mouse models

Mice were housed at the Indiana University School of Medicine (IUSM) animal care facility and were maintained according to USDA standards (12-hr light/dark cycle, food and water ad libitum), per the Guide for the Care and Use of Laboratory Animals (National Institutes of Health, Bethesda, MD).

We used 5xFAD mice which express five human familial Alzheimer’s disease mutations driven by the mouse Thy1 promoter (Oakley et al., 2006) (The Jackson Laboratory [B6SJL-Tg (APPSwFlLon, PSEN1*M146L*L286V)6799Vas, Stock #34840-JAX]). 5xFAD mice as well as its healthy littermate control (C57BL/6J wild-type mice) were aged to 4, 6 and 8 months to model disease progression.

To generate an AD mice model that conditionally inactivate HK2 expression in microglia, we crossed 5xFAD mice expressing cre only in microglia over the CX3CR1 promoter (5xFAD:CX3CR1-creERT2) with mice bearing one or two floxed alleles of HK2 (HK2flox/flox) (Patra et al., 2013). The resulting 5xFAD: CX3CR1-creERT2:HK2flox/wt and 5xFAD: CX3CR1-creERT2:HK2flox/Flox mice (termed 5xFAD:HK2Flox/wt and 5xFAD:HK2Flox/wt respectively) were treated with tamoxifen at 2 months to induce recombination and then aged to 5 months to assess the effect of HK2 haploinsufficiency in disease presentation. 5xFAD: CX3CR1-creERT2:HK2wt/wt mice were equally treated with TAM and used as control. All mice were maintained on a C57BL/6 J background. In this study, both male and female mice were used for all experiments.

#### Human Subjects

Post-mortem brain tissues from subjects with AD and control subjects were provided in the form of frozen blocks by the Brain Resource Center at Johns Hopkins. AD cases consisted of pathologically severe AD, stage V–VI.

#### Dataset analysis

HK2 expression in GSE33000, dataset was analyzed using the online tool GEO2R (https://www.ncbi.nlm.nih.gov/geo/geo2r/).

#### Cell lines and primary cultures

Protocol of primary neonatal microglia cultures was adapted from (Saura et al., 2003). Briefly, brains from 1 to 3 day-old C57BL/6J mouse pups were isolated and chopped. The tissues were then dissociated by mechanical disruption using a nylon mesh and cells were plated in a six-well-plate in advanced DMEM-F12 medium containing 10% FBS, 1% glutamax and 1% penicillin-streptomycin. Every seven days, medium was replenished. After 3 weeks, mixed glial cultures reached confluence and were isolated by mild trypsinization. Briefly, cells were washed with culture medium without FBS and treated with a mixture of trypsin (0.25% without EDTA) and DMEM-F12 medium in a 1:3 ratio. After 15 min incubation, astrocytic cell layer detached and left a layer of microglia attached to the bottom of the culture dish. Microglial cells were maintained in mixed glial cell media for 2 to 3 extra days.

### Method Details

#### Drug treatments

PLX5622 was provided by Plexxikon formulated in AIN-7 diet at 1200 mg/kg. At 4 months of age, either normal rodent diet or PLX5622-containing chow was administered for 28 days. An additional cohort of 4-month-old mice were treated with PLX5622 or control diet for 28 days, then discontinued from PLX5622 feed and treated with a normal rodent diet for an additional 28 days. At 6 months of age, this cohort of mice was euthanized, and molecular analyses were performed. Experiments always used littermate controls.

Lonidamine was purchased from Cayman Chemicals (14640) and stock solution was prepared in DMSO (Sigma D2650). The daily dose of LND used was 50 mg/kg body weight diluted in PBS1x (DMSO 0.05%) and was injected intraperitoneally into 5-month-old 5xFAD mice in 1 ml volume, for 7 consecutive days. The drug as well as the vehicle solution was well tolerated and did not affect the body weight of the mice. For in vitro experiments, 200 μM of LND was added to the media 30 min prior to its co-incubation with fAβ (5 μM). For gene expression analysis, the primary cultures were incubated with the same concentration of LND for 24 hrs. For hexokinase activity, cellular lysates were incubated for 1h with LND or vehicle with was measured using a hexokinase assay kit (Abcam, ab211103) according to manufacturer instructions.

Tamoxifen was obtained from Sigma (T5648). Stock solutions were prepared in vehicle (10% Ethanol (Sigma E7023) + 90% corn oil (Sigma C8267)) and stored protected from light at -20 °C. The daily dose of TAM used was 15 mg/kg body weight and was injected intraperitoneally into 2-month-old 5xFAD mice in 0.1 ml volume, for 5 consecutive days.

#### Spontaneous Alternation Y-Maze Test

Mice were acclimated to the testing room under ambient lighting conditions for 1h. Spontaneous alternation performance was tested using a symmetrical Y-maze. Mice were placed in the center of the Y-maze and the sequence of entries into each arm was recorded via a ceiling-mounted HD camera integrated with behavioral tracking software (ANY-maze. Stoelting, CO, USA). Each mouse was placed and was allowed to explore freely through the maze during an 8-min session. The sequence and total number of arms entered were recorded. Arm entry was complete when the hind paws of the mouse had been completely placed in the arm. Percentage alternation was calculated as the number of triads containing entries into all three arms divided by the maximum possible alternations (the total number of arms entered minus 2) × 100. After behavioral testing, mice were sacrificed for biochemical and histological analysis.

#### Collection of tissue and tissue homogenization

Following treatment, animals were anesthetized and euthanized according to the IUSM Institutional Animal Care and Use Committee-approved procedures. After transcardial perfusion with PBS 1X, mouse brains hemispheres were divided for immunohistochemistry (IHC) studies and homogenization. One hemisphere was fixed by overnight with 4% PFA at 4 °C. Then, brains were cryoprotected in 30% sucrose at 4°C and embedded. Brains were processed on a cryostat as 30 μm free-floating sections. The other hemisphere was homogenized in Tissue homogenization buffer (THB) supplemented with protease inhibitor cocktail. One portion of the homogenate was further sonicated for protein extraction and western blot and ELISA analysis and the other was stored at -80 °C in RNA bee (Amsbio, CS-501B) for RNA extraction and gene expression analysis.

Post-mortem brain tissues from subjects with AD were homogenized in RIPA buffer with a protease inhibitor cocktail (Roche) and a dilution of brain to RIPA of 1:10 (w/v). Samples were then centrifuged at 13,200 rpm for 15 min at 4°C. The supernatants were portioned into aliquots, snap-frozen, and stored at −80°C until analyzed. The RIPA-insoluble pellet was treated with formic acid by mixing samples with 88% formic acid for 1 hr at room temperature (the volume of 88% FA was one-fourth of the volume used for RIPA). Samples were then diluted with distilled water to obtain the same volume used in RIPA and lyophilized for 24 hr. Freeze-dried samples were reconstituted in PBS using the same volume that was originally used for RIPA. Samples were then sonicated for 30 s. Finally, samples were mixed with running buffer, run on a gel, and analyzed by western blot.

#### Immunostaining, image acquisition, and image analysis

For immunostaining, at least three matched brain sections were used. Free-floating sections were washed and permeabilized in 0.1% Triton in PBS (PBST), followed by antigen retrieval using 1x Reveal decloacker (Biocare medical) at 85 °C for 15 min. Sections were blocked in 5% normal donkey serum in PBST for 1 h at room temperature (RT). Primary antibodies were incubated in 5% normal donkey serum in PBST overnight at 4 °C. Sections were washed and visualized using respective species-specific AlexaFluor fluorescent antibodies (diluted 1:1000 in 5% normal donkey serum in PBST for 1 h at RT). Sections were counterstained and mounted onto slides. For thioflavin-S (ThioS) staining (Sigma, T1892), sections were dried at room temperature, rehydrated in PBS, and stained in 0.1% volume ThioS solution for 5 min at RT. Sections were then washed twice for 2 min each in 70% ethanol, washed again in PBS 1X, and then mounted in ProLong gold (Invitrogen, P36930). Images were acquired on a Nikon confocal microscope with z step set at 1 μm of thickness with a 10x air objective for evaluate immunoreactive areas and 60x oil objective for high resolution imaging of plaque associated microglia.

Three brain sections were analyzed for each animal. For each brain section, two regions of interest were imaged with base in their high amyloid levels: subiculum and somatosensory cortex. Images were analyzed using ImageJ (NIH). To determine immunoreactive area, Images were manually thresholded and maximum projections were prepared from z-stacks. Immunoreactive area was determined using the ‘measure’ function and averaged per area to determine the percentage of immunoreactive coverage. Aβ plaque counts were performed using the “analyze particles” function after manual threshold applied as mentioned above, and mask was applied to the images. For all applicable quantifications, physical area in square μm was determined using the ‘measure’ command of ImageJ for each region. Quantification is represented as either percent area coverage or count per square μm.

#### RNA isolation and quantitative, real-time PCR

RNA was extracted from homogenized tissue using PureLink RNA Mini Columns following the manufacturer’s instructions. RNA was reverse transcribed into cDNA using the High-Capacity RNA to cDNA set. Taqman MasterMix and StepOne Plus (Applied Biosystems) was used for qPCR as per the manufacturer’s instructions. For all mRNA analyses, housekeeping gene GAPDH was used. Results are represented as relative fold change in gene expression normalized to the wild-type calibrator. For statistics, the relative delta Ct method was used.

#### Nanostring

The nCounter Analysis System (NanoString Technologies, Seattle, WA, USA) allows for multiplexed digital mRNA profiling without amplification or generation of cDNA (Geiss et al., 2008). Then nCounter Glial profiling panel profile 770 mouse genes across 50+ pathways involved in glial cell biology and quantify the relative abundance of 5 brain cell types and peripheral immune cells. Briefly, 200 ng of RNA was loaded for all samples and hybridized with probes for 16 h at 65 °C. Results obtained from nCounter MAX Analysis System (NanoString Technologies, catalog #NCT-SYST-LS, Seattle WA) were imported to Rosalind Analysis Platform (OnRamp Bioinformatics) for QC verification, normalization, and data statistics using Advanced Analysis (v2.0.115; NanoString Technologies). Probes were only included if the read count was more than 3 standard deviations above background, and probes that had <100 reads for 6 or more samples were removed from analysis. All assays were performed according to manufacturer protocols.

#### Western blot

After sonication, the soluble fraction of brain lysates was obtained by centrifugation for 10 min at 14,000 rpm. The protein samples were measured via Pierce BCA assay (Thermo-Fisher, 23225). For protein identification and relative quantification, 10 to 25 Lg of proteins loaded onto Bolt 4%–12% Bis-Tris plus Protein gels (Thermo-Fisher, NW04122BOX) and run at 200 mV for 40 min. The proteins were then transferred onto PVDF membrane (Millipore, IPVH00010) at 400 mAmp for 90 min on ice. After 1 h of blocking in 5% BSA (in PBS 1X) the primary antibodies were then applied overnight in a blocking buffer at 4LC. The HRP-conjugated secondary antibodies were all incubated for 1h at RT at the proper dilution. The signal was developed using Immobilon western Chemiluminescent HRP Substrate (Millipore, WBKLS0500). Images were acquired with Amersham imager 600 (General-Electrics Healthcare) and protein quantification was performed by measuring the optical density of the specific bands with image J software (NIH). Samples from all experimental groups were loaded on each experiment and normalized to the respective control. For statistical analysis, normalized data across independent experiments were used together.

#### Cytokine Panel Assay

Mouse brain samples were assayed in duplicate using the MSD Proinflammatory Panel I (K15048D; MesoScale Discovery, Gaithersburg, MD, United States), a highly sensitive multiplex enzyme-linked immunosorbent assay (ELISA). This panel quantifies the following 10 proinflammatory cytokines in a single small sample volume (25 μL) of supernatant using an electrochemiluminescent detection method (MSD): interferon γ (IFN-γ), interleukin (IL)-1β, IL-2, IL-4, IL-6, IL-8, IL-10, IL-12p70, IL-13, and tumor necrosis factor α (TNFα). The mean intra-assay coefficient for each cytokine was <8.5%, based on the cytokine standards. Any value below the lower limit of detection (LLOD) for the cytokine assay was replaced with 1/2 LLOD of the assay for statistical analysis.

#### Flow cytometry

For in vivo phagocytosis evaluation, 3 hr before brain harvest, 5-month old mice were i.p. injected with a blood–brain barrier-penetrating Aβ dye, methoxy-XO4. If microglia are actively phagocytosing amyloid plaques, methoxy-XO4 will be internalized together with Aβ. Mice were perfused, brains removed, chopped, and digested using the Macs Neural tissue Dissociation kit (Miltenyl Biotec) and subsequent percoll gradients (30% percoll [GE healthcare], 10% fetal bovine serum) was used to purify myeloid cells and incubated to antibodies against Cd11b and CD45. Later, stained samples were fixed and permeabilized using the BD Cytofix/Cytoperm kit. Intracellular staining of HK2 was done by using a rabbit monoclonal antibody (Abcam) and an anti-rabbit conjugated secondary antibody. To evaluate microglial mitochondrial membrane potential (ΔΨM), we used Mitotracker deep red (DR). Samples were acquired on BD FACS Canto II (BD Bioscience) flow cytometer. Raw data were analyzed with FlowJo v10 (BD Bioscience). Cells were gated on Cd11b+ and Cd45low to assess the level of HK2 expression, Aβ phagocytosis and ΔΨM as the geometric mean fluorescence intensity (gMFI).

#### Lonidamine quantification in mice brain and plasma

Brains and blood of Lonidamine treated mice were collected, weighed, crushed to a powder with mortar and pestle under liquid nitrogen, and powdered brains were stored at -80°C until analysis. A method to quantify Lonidamine from mouse brain and plasma has been developed using temazepam (TMP, Sigma) as the internal standard, liquid-liquid extraction, and HPLC-MS/MS (Sciex 5500 QTRAP). The mass spectrometer utilizes an electrospray ionization probe run in positive mode. The multiple reaction monitoring (MRM) Q1/Q3 (m/z) transitions for Lonidamine was 322.1/304.0 and temazepam is 301.1/255.0. The lower limit of quantification using 20 µL of mouse plasma was 3 ng/mL and for whole mouse brain was 2.4 ng/sample. The extraction uses phosphate buffer, pH 7.4, and methyl tertiary butyl ether. The mobile phase was delivered via gradient using acetonitrile and 0.1% formic acid on a Restek Ultra C18 50X4.6mm 5µm column.

#### Measurements of ATP levels, HK activity and MTT assay

Primary microglia were seeded in a 96 well plate and after 24 hrs, cells were homogenized, and ATP, MTT (cell viability) and HK activity were measured using Luminescent ATP Detection Assay Kit (Abcam, ab113849), Colorimetric hexokinase activity assay kit (Abcam, ab136957) and CellTiter 96 Non-Radioactive Cell Proliferation Assay.

#### Statistics and study analysis

Statistical tests, along with the number of animals analyzed, are indicated in the figure legends. Comparisons between two groups were conducted with the unpaired, two-tailed student’s t -tests, and comparisons between multiple groups were conducted with the one-way ANOVA with Tukey’s post-hoc multiple comparison test using Prism 9 statistical software (GraphPad, San Diego, CA, United States). Error bars represent the standard error of the mean (SEM). Statistical significance was met when the p -value was less than 0.05.

## Acknowledgments

The authors would like to thank to Dr. Nissim Hay from the Departments of Biochemistry and Molecular Genetics at the University of Illinois College of Medicine, for his generous gift of the HK2 floxed mice. We appreciate the collaboration of Dr. Jheel Patel for her help in the behavioral testing. Lonidamine quantification was provided by the Clinical Pharmacology Analytical Core (CPAC) at Indiana University School of Medicine, supported by the IU Simon Comprehensive Cancer Center Support Grant P30 CA082709.

This work was supported by grant from National Institutes of Health (NIH) (to G. E. L., RF1AG068400). J.F.C. was supported by Eli Lilly-Stark Neuroscience Research grant (EPAR1536) and BrightFocus Foundation (A20201166F).

## Author Contributions

J.F.C. and G. E. L. designed the experiments and wrote the manuscript. J.F.C. and C.M.R. performed the experiments. P.B.L. performed the Elisa experiments B. T. C. performed the microglial depletion experiments and provided the samples for analysis of this project. S. S. P. performed the flow cytometry experiments. C. L. R., N. J. and P. M. provided and analyzed the human samples. All the authors assisted in editing the manuscript.

## Supplementary figure legends

**Fig S1.**
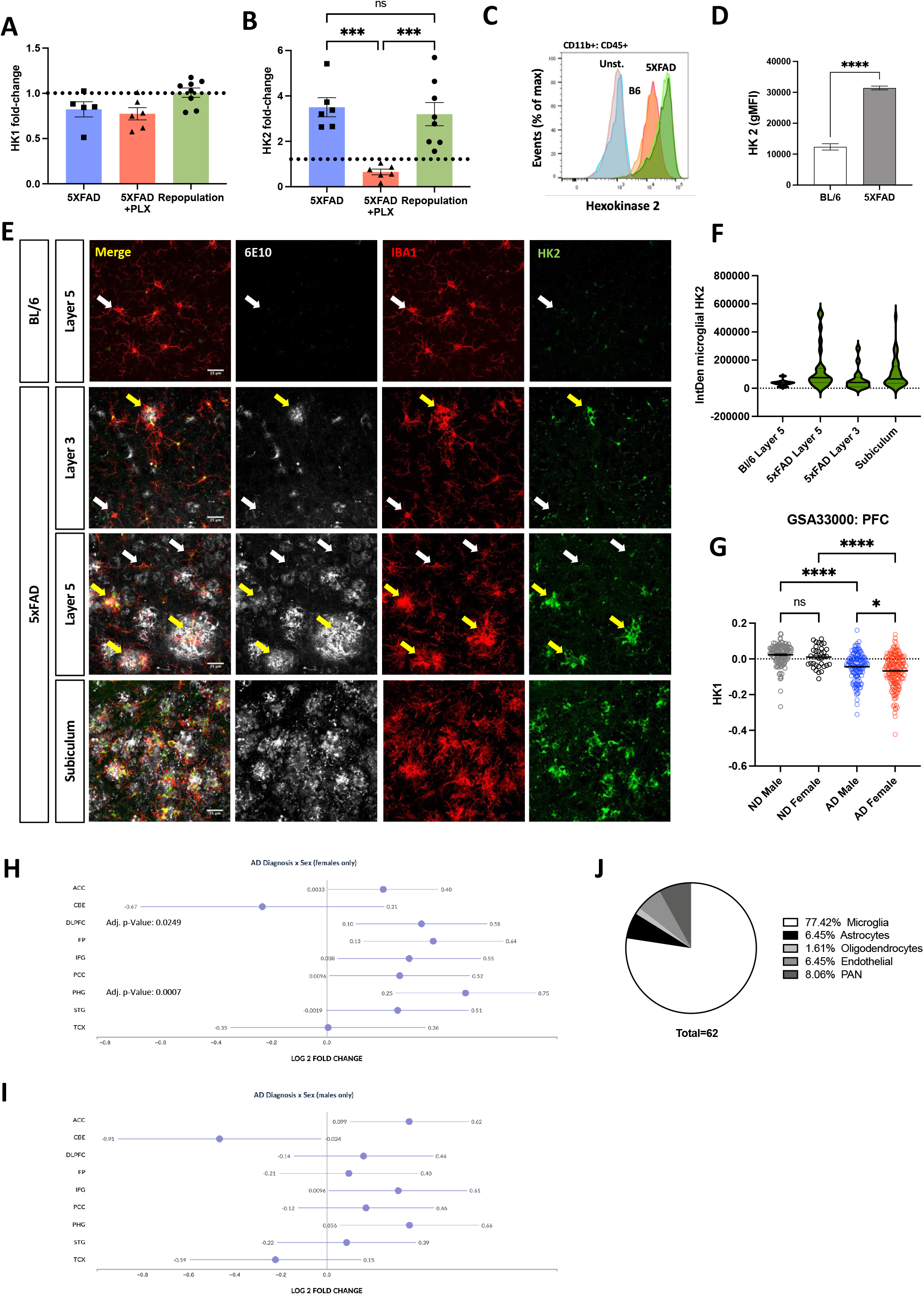
Microglial HK2 upregulation is associated to the presence of Aβ plaques. **(A and B)** qPCR analysis of HK1 and HK2 expression in the cortex of 4-mo-old 5xFAD mice depleted of microglia by PLX treatment and after 28 days of microglial repopulation ***p<0.001, ****p< 0,0001 (n=4-8 per group, One-way ANOVA followed by Tukey’s post-hoc test). **(C)** Representative histograms comparing unstained and HK2 stained 5xFAD (green) and Bl/6 (orange) CD11b^+^, CD45^+^ microglia. **(D)** Quantification of the HK2 geometric mean fluorescence intensity (gMFI) of CD11b^+^, CD45^+^ microglia shown in **E**. ****p< 0,0001 (n=3, Unpaired t test). **(E)** IHC of HK2 expression (green) in Iba1+ microglia (red) across different layers of the cortex and subiculum of the 5xFAD mice and it healthy control. **Upper panel.** IHC of Bl/6 layer 5 cortex is shown as an example of HK2 expression in homeostatic microglia (white arrows). **Lower panels.** IHC of 5xFAD subiculum and cortical layer 3 and 5. Aβ-clustered microglia shown increased levels of HK2 (yellow arrows) in comparation to the ramified parenchymal microglia. (F) Quantification of the integrated density of HK2 in Iba1+ cells. After binarization of Iba1 immunoreactive signal, the mean fluorescence intensity of HK2 was calculated and multiplied by the area of Iba1. *p<0.05 (n=8-9 per group, Unpaired t test). (G) HK1 expression obtained from a human transcriptomic dataset of prefrontal cortex tissue (PFC) of 157 nondemented controls and 310 AD patients (GSE33000). Data segregated by sex reveals a significant difference in the down-regulated levels of HK1 between woman and men. **(H and I)** AMP-AD consortium analysis of the differential expression of HK2 across brain regions in women (H) and men (I) AD donors. Forest plot indicates the estimate of the log fold change with standard errors across different brain regions. **(J)** Cellular expression of genes co-expressed with HK2 described in Fig. 1J. The percentage of cellular specificity for each gene was evaluated using an RNA-Seq database of cell types isolated from mouse and human brain. https://www.brainrnaseq.org

**Fig S2.**
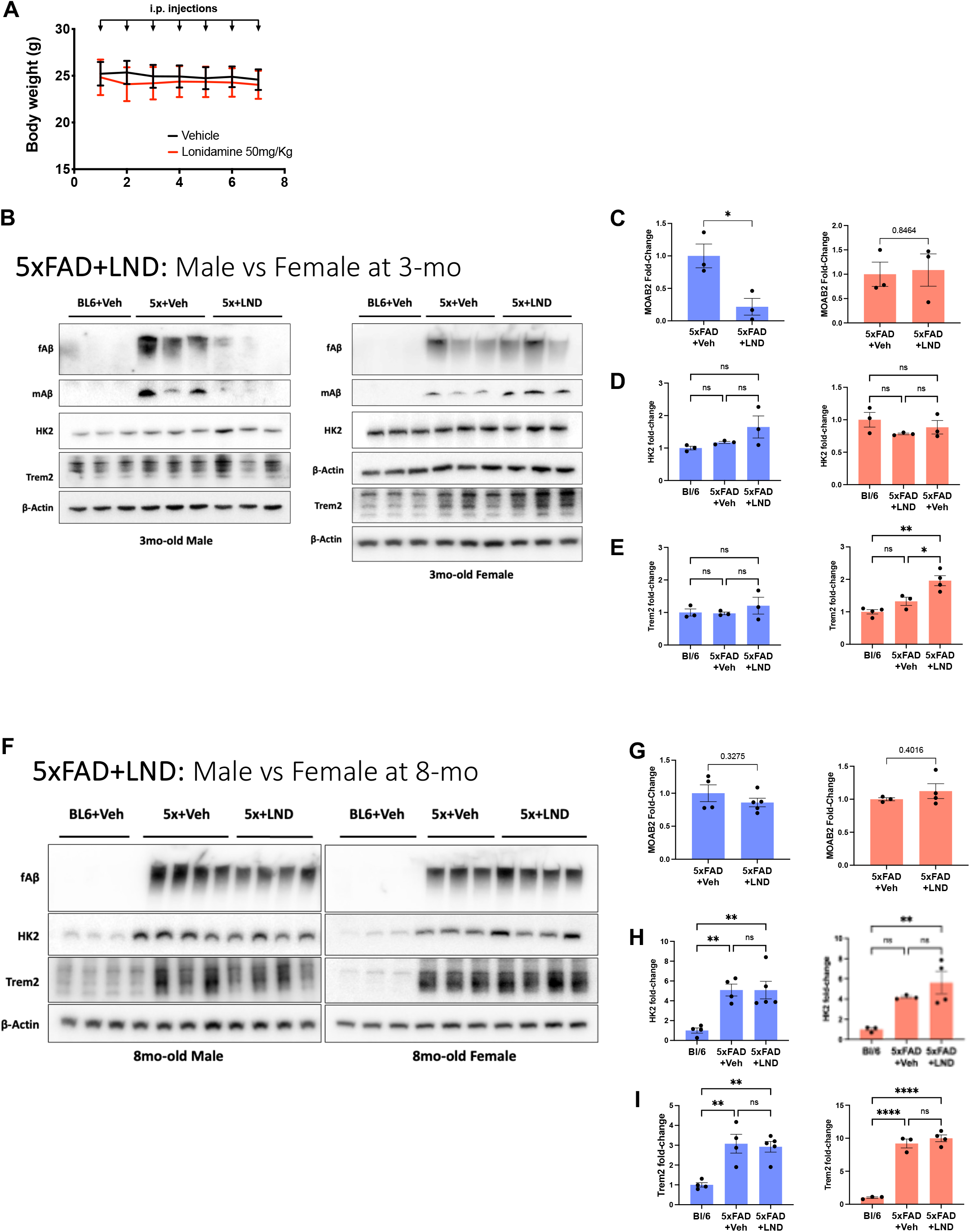
Sex biased response to LND was not dependent of pathology severity. **(A)** Administration of HK2 pharmacological inhibitor, Lonidamine (LND) did not affected the body weight of treated mice during the 7 days of i.p. administration. **(B-E)** Western blot and quantification of proteins from soluble fraction of cortical lysates of BL/6 and 5xFAD male mice treated with vehicle or LND at 3 months of age (B) Representative immunoblots of fibrillar and monomeric Aβ detected with MOAB-2 antibody, microglial proteins HK2, and Trem2. Left immunoblot show the effect of LND in male cortex and right immunoblot in female (C-E) Immunoreactive bands were quantified, normalized to β-actin and expressed as the percentage of 5xFAD +veh mice in the case of MOAB-2. *p< 0.05 (n=3, Unpaired t-test). For the other proteins the values are expressed as the percentage of BL/6 +veh mice. (*p< 0.05, **p< 0.01. n=3, One-way ANOVA, Tukey posttest). **(F-I)** Western blot and quantification of proteins from soluble fraction of cortical lysates of BL/6 and 5xFAD male mice treated with vehicle or LND at 8 months of age as described before. For MOAB-2, n=3-5, Unpaired t-test. For the other proteins the values are expressed as the percentage of BL/6 +veh mice. **p< 0.01, ****p< 0,0001. (n=3-5, One-way ANOVA, Tukey posttest).

**Fig S3.**
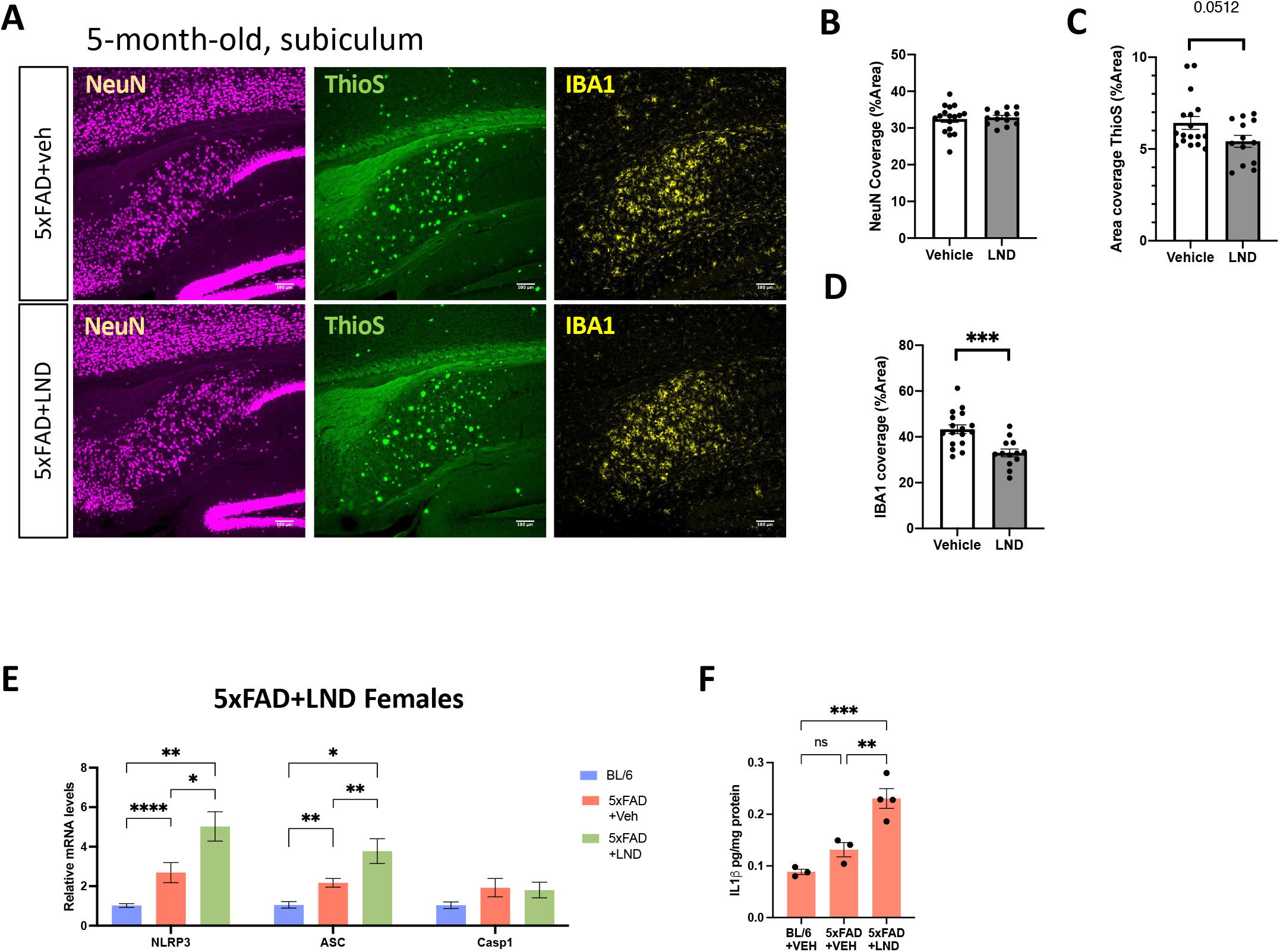
Pharmacological inhibition of HK2 with LND fails to improve pathology in a context dependent manner. **(A)**Immunohistochemical analysis of the subiculum area of 5-mo-old 5xFAD male mice treated with LND or vehicle. Amyloid plaques were stained with ThioS (green), neurons with NeuN (purple) and microglia with Iba1 (yellow). Scale bar 100 μm. **(B-D)** Quantification of subiculum area fraction of NeuN (B), ThioS (C) and Iba1 • (D). ***p<0.001 (n=8-10 per group, Unpaired t-test) **(E)** qPCR analysis of elements of the inflammasome in females Bl/6, 5xFAD and 5xFAD treated with LND. *p< 0.05, **p< 0.01, ****p< 0,0001 (n=3-4, One-way ANOVA, tukey posttest). **(F)** ELISA analysis of IL-1β levels in LND treated females 5xFAD mice. **p< 0.01, ***p< 0,001 (n=3-4, One-way ANOVA, tukey posttest)

## References

1. Sasaki, A., Yamaguchi, H., Ogawa, A., Sugihara, S. & Nakazato, Y. Microglial activation in early stages of amyloid beta protein deposition. Acta Neuropathol. 94, 316–22 (1997).

2. Yuan, P. et al. TREM2 Haplodeficiency in Mice and Humans Impairs the Microglia Barrier Function Leading to Decreased Amyloid Compaction and Severe Axonal Dystrophy. Neuron 90, 724–39 (2016).

3. d’Errico, P., et al. Microglia contribute to the propagation of Aβ into unaffected brain tissue. Nat. Neurosci. 25, 20–25 (2022).

4. Wang, C. et al. Microglial NF-κB drives tau spreading and toxicity in a mouse model of tauopathy. Nat. Commun. 13, 1969 (2022).

5. Jain, N., Lewis, C. A., Ulrich, J. D. & Holtzman, D. M. Chronic TREM2 activation exacerbates Aβ-associated tau seeding and spreading. J. Exp. Med. 220, (2023).

6. Ghosh, S., Castillo, E., Frias, E. S. & Swanson, R. A. Bioenergetic regulation of microglia. Glia (2017) doi:10.1002/glia.23271.

7. Loftus, R. M. & Finlay, D. K. Immunometabolism: Cellular Metabolism Turns Immune Regulator. J. Biol. Chem. 291, 1–10 (2016).

8. Paolicelli, R. C. & Angiari, S. Microglia immunometabolism: From metabolic disorders to single cell metabolism. Semin. Cell Dev. Biol. 94, 129–137 (2019).

9. Gimeno-Bayón, J., López-López, A., Rodríguez, M. J. & Mahy, N. Glucose pathways adaptation supports acquisition of activated microglia phenotype. J. Neurosci. Res. 92, 723–731 (2014).

10. Fairley, L. H., Wong, J. H. & Barron, A. M. Mitochondrial Regulation of Microglial Immunometabolism in Alzheimer’s Disease. Front. Immunol. 12, 624538 (2021).

11. Codocedo, J. F. & Landreth, G. E. The intersection of metabolism and inflammation is governed by the intracellular topology of hexokinases and the metabolic fate of glucose. Immunometabolism (Cobham (Surrey, England)) 4, e00011 (2022).

12. Granchi, C. & Minutolo, F. Anticancer Agents That Counteract Tumor Glycolysis. ChemMedChem 7, 1318–1350 (2012).

13. Baik, S. H., et al. Hexokinase dissociation from mitochondria promotes oligomerization of VDAC that facilitates NLRP3 inflammasome assembly and activation. Sci. Immunol. 8, eade7652 (2023).

14. Leng, L. et al. Microglial hexokinase 2 deficiency increases ATP generation through lipid metabolism leading to β-amyloid clearance. Nat. Metab. 4, 1287– 1305 (2022).

15. Hu, Y. et al. Dual roles of hexokinase 2 in shaping microglial function by gating glycolytic flux and mitochondrial activity. Nat. Metab. 4, 1756–1774 (2022).

16. Cheng, J. et al. Early glycolytic reprogramming controls microglial inflammatory activation. J. Neuroinflammation 18, 129 (2021).

17. Wolf, A. J. et al. Hexokinase Is an Innate Immune Receptor for the Detection of Bacterial Peptidoglycan. Cell 166, 624–636 (2016).

18. Sil, A. et al. Sex Differences in Behavior and Molecular Pathology in the 5XFAD Model. J. Alzheimers. Dis. 85, 755–778 (2022).

19. Casali, B. T., MacPherson, K. P., Reed-Geaghan, E. G. & Landreth, G. E. Microglia depletion rapidly and reversibly alters amyloid pathology by modification of plaque compaction and morphologies. Neurobiol. Dis. 142, 104956 (2020).

20. Keren-Shaul, H. et al. A Unique Microglia Type Associated with Restricting Development of Alzheimer’s Disease. Cell 169, 1276–1290.e17 (2017).

21. Krasemann, S. et al. The TREM2-APOE Pathway Drives the Transcriptional Phenotype of Dysfunctional Microglia in Neurodegenerative Diseases. Immunity 47, 566–581.e9 (2017).

22. Morgan, B. P. Complement in the pathogenesis of Alzheimer’s disease. Semin. Immunopathol. 40, 113–124 (2018).

23. De Schepper, S. et al. Perivascular cells induce microglial phagocytic states and synaptic engulfment via SPP1 in mouse models of Alzheimer’s disease. Nat. Neurosci. 26, 406–415 (2023).

24. Liu, S. et al. TLR2 is a primary receptor for Alzheimer’s amyloid β peptide to trigger neuroinflammatory activation. J. Immunol. 188, 1098–107 (2012).

25. Guillot-Sestier, M.-V. et al. Microglial metabolism is a pivotal factor in sexual dimorphism in Alzheimer’s disease. Commun. Biol. 4, 711 (2021).

26. Lynch, M. A. Exploring Sex-Related Differences in Microglia May Be a Game-Changer in Precision Medicine. Front. Aging Neurosci. 14, 868448 (2022).

27. Patra, K. C. et al. Hexokinase 2 is required for tumor initiation and maintenance and its systemic deletion is therapeutic in mouse models of cancer. Cancer Cell 24, 213–228 (2013).

28. Oakley, H. et al. Intraneuronal beta-amyloid aggregates, neurodegeneration, and neuron loss in transgenic mice with five familial Alzheimer’s disease mutations: potential factors in amyloid plaque formation. J. Neurosci. 26, 10129–40 (2006).

29. Wolf, A., Bauer, B., Abner, E. L., Ashkenazy-Frolinger, T. & Hartz, A. M. S. A Comprehensive Behavioral Test Battery to Assess Learning and Memory in 129S6/Tg2576 Mice. PLoS One 11, e0147733 (2016).

30. Garcia, S. N., Guedes, R. C. & Marques, M. M. Unlocking the potential of HK2 in cancer metabolism and therapeutics. Curr. Med. Chem. (2018) doi:10.2174/0929867326666181213092652.

31. Cervantes-Madrid, D., Romero, Y. & Dueñas-González, A. Reviving Lonidamine and 6-Diazo-5-oxo-L-norleucine to Be Used in Combination for Metabolic Cancer Therapy. Biomed Res. Int. 2015, 690492 (2015).

32. Huang, Y. et al. The Potential of Lonidamine in Combination with Chemotherapy and Physical Therapy in Cancer Treatment. Cancers (Basel). 12, 3332 (2020).

33. Feng, W. et al. Microglia prevent beta-amyloid plaque formation in the early stage of an Alzheimer’s disease mouse model with suppression of glymphatic clearance. Alzheimers. Res. Ther. 12, 125 (2020).

34. Thawkar, B. S. & Kaur, G. Inhibitors of NF-κB and P2X7/NLRP3/Caspase 1 pathway in microglia: Novel therapeutic opportunities in neuroinflammation induced early-stage Alzheimer’s disease. J. Neuroimmunol. 326, 62–74 (2019).

35. Oblak, A. L. et al. Comprehensive Evaluation of the 5XFAD Mouse Model for Preclinical Testing Applications: A MODEL-AD Study. Front. Aging Neurosci. 13, 713726 (2021).

36. Bianco, A., Antonacci, Y. & Liguori, M. Sex and Gender Differences in Neurodegenerative Diseases: Challenges for Therapeutic Opportunities. Int. J. Mol. Sci. 24, (2023).

37. Zucker, I. & Prendergast, B. J. Sex differences in pharmacokinetics predict adverse drug reactions in women. Biol. Sex Differ. 11, 32 (2020).

38. Wang, P., Henning, S. M. & Heber, D. Limitations of MTT and MTS-based assays for measurement of antiproliferative activity of green tea polyphenols. PLoS One 5, e10202 (2010).

39. Gao, C., Jiang, J., Tan, Y. & Chen, S. Microglia in neurodegenerative diseases: mechanism and potential therapeutic targets. Signal Transduct. Target. Ther. 8, 359 (2023).

40. Venegas, C. et al. Microglia-derived ASC specks cross-seed amyloid-β in Alzheimer’s disease. Nature 552, 355–361 (2017).

41. Zipp, F., Bittner, S. & Schafer, D. P. Cytokines as emerging regulators of central nervous system synapses. Immunity 56, 914–925 (2023).

42. Demarquoy, J. & Le Borgne, F. Crosstalk between mitochondria and peroxisomes. World J. Biol. Chem. 6, 301–9 (2015).

43. Shen, Y. et al. Bioenergetic state regulates innate inflammatory responses through the transcriptional co-repressor CtBP. Nat. Commun. 8, 624 (2017).

44. Lin, J. et al. Glycolytic enzyme HK2 promotes PD-L1 expression and breast cancer cell immune evasion. Front. Immunol. 14, 1189953 (2023).

45. Guo, D. et al. Aerobic glycolysis promotes tumor immune evasion by hexokinase2-mediated phosphorylation of IκBα. Cell Metab. 34, 1312–1324.e6 (2022).

46. Baik, S. H. et al. A Breakdown in Metabolic Reprogramming Causes Microglia Dysfunction in Alzheimer’s Disease. Cell Metab. 30, 493–507.e6 (2019).

47. Lynch, M. A. Can the emerging field of immunometabolism provide insights into neuroinflammation? Prog. Neurobiol. 184, 101719 (2020).

48. Seki, S. M. & Gaultier, A. Exploring Non-Metabolic Functions of Glycolytic Enzymes in Immunity. Front. Immunol. 8, 1549 (2017).

49. Vilalta, A. & Brown, G. C. Deoxyglucose prevents neurodegeneration in culture by eliminating microglia. J. Neuroinflammation 11, 58 (2014).

50. Hinrichsen, F. et al. Microbial regulation of hexokinase 2 links mitochondrial metabolism and cell death in colitis. Cell Metab. 33, 2355–2366.e8 (2021).

51. Sayed, F. A. et al. Differential effects of partial and complete loss of TREM2 on microglial injury response and tauopathy. Proc. Natl. Acad. Sci. U. S. A. 115, 10172–10177 (2018).

52. Andreone, B. J. et al. Alzheimer’s-associated PLCγ2 is a signaling node required for both TREM2 function and the inflammatory response in human microglia. Nat. Neurosci. 23, 927–938 (2020).

53. Waters, A., Society for Women’s Health Research Alzheimer’s Disease Network & Laitner, M. H. Biological sex differences in Alzheimer’s preclinical research: A call to action. Alzheimer’s Dement. (New York, N. Y.) 7, e12111 (2021).

